# A dopamine-induced gene expression signature regulates neuronal function and cocaine response

**DOI:** 10.1101/781872

**Authors:** Katherine E. Savell, Morgan E. Zipperly, Jennifer J. Tuscher, Corey G. Duke, Robert A. Phillips, Allison J. Bauman, Saakshi Thukral, Faraz A. Sultan, Nicholas A. Goska, Lara Ianov, Jeremy J. Day

## Abstract

Drug addiction is a worldwide health problem, with overdose rates of both psychostimulants and opioids currently on the rise in many developed countries. Drugs of abuse elevate dopamine levels in the nucleus accumbens (NAc) and alter transcriptional programs believed to promote long-lasting synaptic and behavioral adaptations. However, even with well-studied drugs such as cocaine, drug-induced transcriptional responses remain poorly understood due to the cellular heterogeneity of the NAc and complex drug actions via multiple neurotransmitter systems. Here, we leveraged high-throughput single-nucleus RNA-sequencing to create a comprehensive molecular atlas of cell subtypes in the NAc, defining both sex-specific and cell type-specific responses to acute cocaine experience in a rat model system. Using this transcriptional map, we identified specific neuronal subpopulations that are activated by cocaine, and defined an immediate early gene expression program that is upregulated following cocaine experience *in vivo* and dopamine (DA) receptor activation *in vitro*. To characterize the neuronal response to this DA-mediated gene expression signature, we engineered a large-scale CRISPR/dCas9 activation strategy to recreate this program. Multiplexed induction of this gene program initiated a secondary synapse-centric transcriptional profile, altered striatal physiology *in vitro*, and enhanced cocaine sensitization *in vivo*. Taken together, these results define the genome-wide transcriptional response to cocaine with cellular precision, and demonstrate that drug-responsive gene programs are sufficient to initiate both physiological and behavioral adaptations to drugs of abuse.

---

NEARLY 5 MILLION Americans reported cocaine use in 2017, and recent increases in cocaine-related drug overdoses present significant public health challenges (*1*). A hallmark trait of drugs of abuse is the acute elevation of dopamine (DA) in the nucleus accumbens (NAc), a central integrator of the reward circuit (*2-4*). Abused drugs produce increases in DA that are greater in both concentration and duration than natural rewards (*2, 5, 6*), and this signaling is hypothesized to underlie maladaptive reinforcement after repeated drug use (*7*). Exposure to drugs of abuse results in significant transcriptional and epigenetic reorganization in the NAc (*8-13*), initiating synaptic and behavioral plasticity associated with the transition to drug addiction (*7, 14, 15*). However, even with well-studied drugs such as cocaine, drug-induced transcriptional responses remain poorly understood. This is in part due to the cellular heterogeneity of the NAc, which is a diverse structure containing multiple neuronal and non-neuronal subpopulations and complex neuronal circuitry. Additionally, many drugs of abuse engage multiple neurotransmitter systems in the NAc (*16-22*), and the specific contributions of DA-dependent transcriptional programs to neuronal physiology and behavior is not clear. Further, although drug experience leads to large scale transcriptional changes in the NAc, previous studies have focused on individual drug-responsive genes instead of coordinated gene programs. Here, we sought to define DA-driven gene expression signatures with single-cell precision and understand the molecular and physiological consequences of this gene program.

To map the transcriptional landscape and cocaine response of the NAc with single-cell resolution, we performed single-nucleus RNA-sequencing (snRNA-seq) on 15,631 NAc cells from both male and female rats after acute exposure to cocaine (**Fig. 1; Supplementary Fig. 1**), using a cocaine dose previously reported to result in enduring synaptic changes in the NAc (*15*). From a merged dataset containing transcriptomes from all cells, unsupervised dimensionality reduction approaches identified discrete NAc neuronal clusters harboring known markers of *Drd1* DA receptor positive and *Drd2* DA receptor positive medium spiny neurons (Drd1-MSNs and Drd2-MSNs, respectively), as well as previously identified (*23, 24*) transcriptionally-defined cell classes including somatostatin (*Sst*) positive interneurons, microglia, astrocytes, polydendrocytes, and oligodendrocytes (**Fig. 1b-c** and **Supplementary Fig. 2**). Notably, this clustering also revealed potentially novel MSN classes marked by expression of *Drd3* (a D2 family DA receptor) and *Grm8* (a metabotropic glutamate receptor). While Drd3-MSN and Grm8-MSN clusters were relatively depleted in the classic MSN marker *Ppp1r1b* (the gene encoding DARPP-32 protein) (*25*), each cluster exhibited high expression of *Foxp2* and *Bcl11b*, genes strongly linked to MSN differentiation and function (**Supplementary Fig. 2**) (*26-28*). Moreover, whereas cell fractions in nearly all clusters were evenly distributed between treatment group (saline and cocaine) and sex, 91% of Drd3-MSNs were found in male animals, indicative of a significant sex bias in this subpopulation (**Supplementary Fig. 3**).

**Figure 1.**
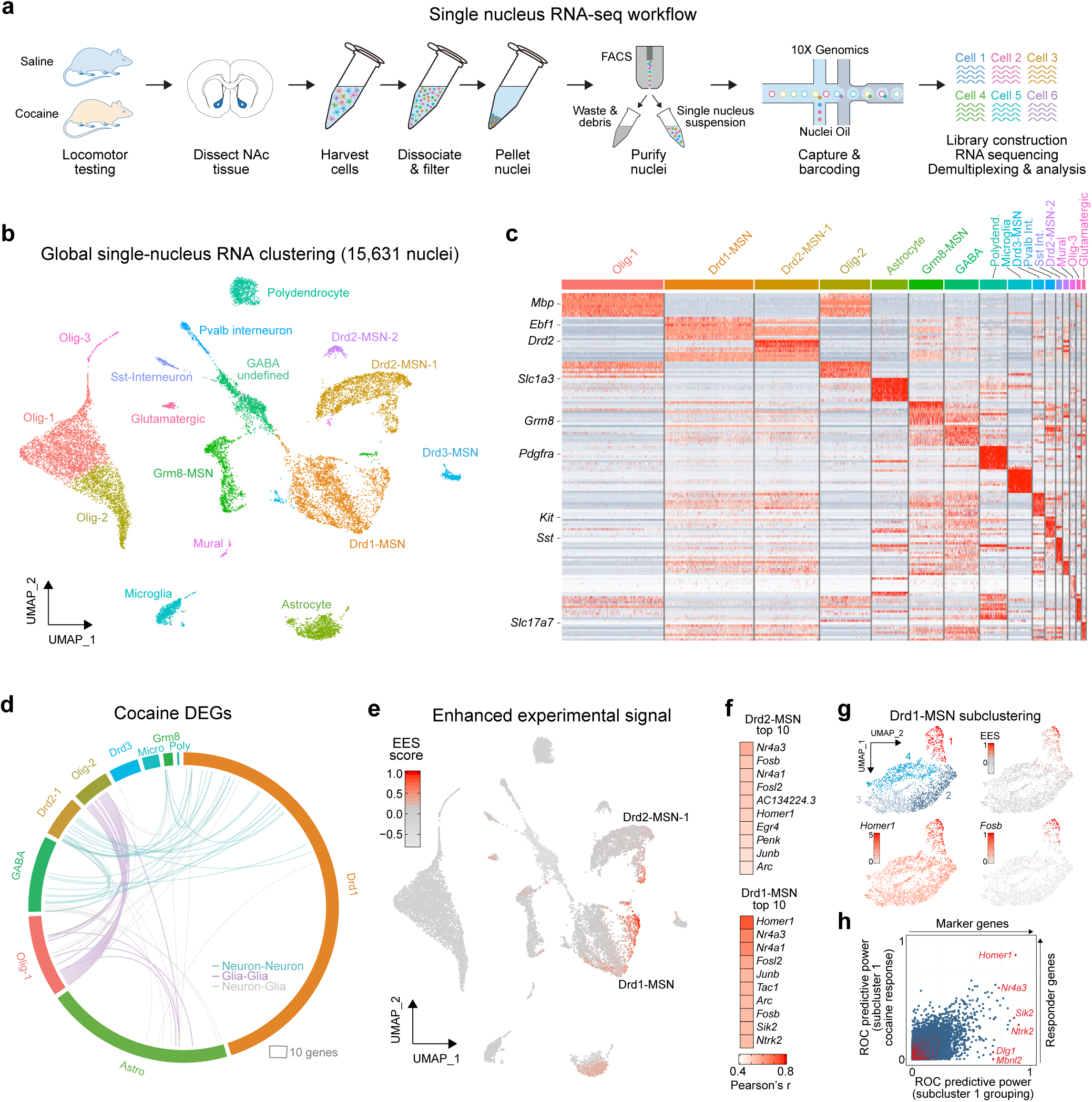
Single-nucleus RNA-seq reveals cell-specific transcriptional response to cocaine. **a**, Single-nucleus RNA-seq workflow. Male and female adult rats (n = 4/sex) received saline or 20mg/kg cocaine (intraperitoneal injection) and underwent locomotor testing prior to tissue harvesting, nucleus accumbens dissection, nuclei purification, and single nuclei sequencing on the 10X Genomics platform. **b**, Global clustering across experimental treatment and sex for 15,631 individual NAc nuclei identifies all major cell classes of the rat NAc, including MSNs expressing *Drd1* and *Drd2* mRNA. **c**, Heatmap of cell-specific marker genes across all clusters (see Supplementary Figure 2 for additional details). **d**, Circos plot of cocaine DEGs by cluster (outer rim, ordered clockwise by adjusted *p* value within each cluster) and coherent changes between clusters (internal arcs). Neuron-neuron, glia-glia, and neuron-glia coherent DEGs are reflected in teal, purple, and gray, respectively. **e**, Unbiased detection of cocaine-activated cell clusters with MELD pipeline identifies enhanced experimental signal (EES) in distinct Drd1-MSN and Drd2-MSN neuronal clusters. **f**, Top 10 gene-EES correlations for Drd1-MSN and Drd2-MSN clusters. **g**, Subclustering of Drd1-MSNs highlights specific IEG induction and EES signal in a single subpopulation (subcluster 1). UMAPs show only Drd1-MSNs identified in panel B. **h**, Receiver operating characteristic (ROC) analysis on subcluster 1 identifies “marker” genes that predict membership in that cluster as compared to all other Drd1-MSN subclusters (x-axis), as well as “responder” genes that predict cocaine treatment classification (i.e., are altered by cocaine).

To identify cocaine-activated cell clusters, we pursued two parallel strategies. First, we performed cluster-specific identification of differentially expressed genes (DEGs) in saline versus cocaine conditions (collapsing across sex; **Fig. 1d; Supplementary Table 1**). This analysis revealed robust transcriptional response in Drd1-MSNs, which contained more DEGs (232) than any other cluster. Drd1-MSN DEGs were enriched in CREB binding motifs and genes involved in MAP kinase pathways, regulation of synaptic signaling, behavior, and cognition (**Supplementary Fig. 4, Supplementary Table 2**). Drd1-MSN DEGs also exhibited overlap with DEGs arising from Drd2-MSNs (**Fig. 1d**), suggesting the induction of common transcriptional pathways in these clusters. We next used an unbiased graphical signal processing approach to stratify cellular clusters based on condition-specific gene signatures (*29*), termed the “enhanced experimental signal”, or EES (**Fig. 1e; Supplementary Table 3**). This method identified two unique cocaine-responsive subclusters – one from the Drd1-MSN parent cluster and another from Drd2-MSN-1 parent cluster. Both high-EES cell clusters exhibited expression of key immediate early genes (IEGs; e.g., *Fosb, Junb*, and *Nr4a1*) in the cocaine condition, but little to no expression of these genes in the saline condition (**Supplementary Fig. 4**). Given that these genes are commonly used markers for neuronal activity (*18, 30*), these results suggest that cocaine activates select ensembles but not the majority of Drd1- or Drd2-MSNs. However, this analysis also revealed correlations between EES and expression of several genes central to neuropeptide signaling in MSNs (*31*), including the substance P precursor *Tac1* (in Drd1-MSNs) and the enkephalin precursor *Penk* (in Drd2-MSNs; **Fig. 1f**).

To determine whether genetic sex contributed to cocaine responses in each cluster, we recalculated EES-gene correlations independently for each sex (**Supplementary Fig. 5, Supplementary Table 4**). Male and female transcriptional responses were positively correlated in both Drd1-MSNs and Drd2-MSNs, and in both cases key IEGs associated with high EES scores in our sex-combined analysis also drove EES scores when sex was considered separately. In contrast, this analysis revealed surprising divergence in astrocytes. For both sexes, numerous astrocyte-responsive genes exhibited strong correlations with EES. However, male and female responses were negatively correlated for this cluster, demonstrating the presence of distinct, non-overlapping, and sex-specific astrocytic transcriptional responses to cocaine.

To further investigate novel upstream interactions that may contribute to cocaine response likelihood within a cell type, we restricted our analysis to Drd1-MSNs and repeated dimensionality reduction mapping approaches (**Fig. 1g**). This analysis identified 4 unique subclusters of Drd1-MSNs, only one of which (subcluster 1) exhibited robust cocaine responsivity. Using this subcluster map, we employed receiver operator characteristic (ROC) analysis to estimate the success of every individual gene in binary classification of a cell in 2 steps. First, we identified genes that marked cells found in the cocaine-activated subcluster (subcluster 1) as compared to cells in all other Drd1-MSN subclusters (termed “marker” genes). Next, we identified genes that distinguished cocaine-treated cells from saline-treated cells in this cocaine-activated subcluster (termed “responder” genes). Intriguingly, while genes such as the synaptic protein *Homer1* served as both marker and responder genes, other genes such as *Dlg1* exhibited high predictive power in marking the cocaine-activated subcluster without exhibiting an altered response to cocaine (**Supplementary Table 5**). In contrast, the *Drd1* receptor itself exhibited almost no predictive power as a marker gene (ROC power = 0.138). Together, these results highlight a potential contribution of multiple genes in determining which Drd1-MSNs are capable of being activated by cocaine.

The activation of specific neuronal ensembles by cocaine may result from circuit-based mechanisms (e.g., differential projections from input structures), or could potentially arise from cell-autonomous mechanisms in response to DA signaling (*7, 17, 32-34*). However, drugs of abuse target many distinct receptor classes in a variety of neuronal and non-neuronal cell types (*20, 21*), and these complex drug actions make identification of DA-induced gene expression programs difficult using *in vivo* models. Therefore, we took advantage of a well-studied and controllable rat primary striatal neuron culture system (*35, 36*) to enable comprehensive and specific identification of DA-dependent gene expression programs in MSNs. Using this system, we first identified a core signature of the transcriptional response to DA receptor activation by performing deep RNA-seq on bulk striatal neuronal cultures treated with 1µM DA for 1hr, a treatment that closely models concentrations and temporal duration of DA increases found *in vivo* after acute cocaine exposure (*2, 3, 6*). This approach identified 103 DEGs (**Fig. 2a-b, Supplemental Table 6**) following DA treatment, with the majority of DEGs (100) being upregulated versus vehicle control samples. Gene ontology analysis revealed that genes upregulated by DA receptor stimulation are primarily associated with transcriptional mechanisms and processes important for neuronal function such as regulation of synaptic plasticity (**Fig. 2c; Supplementary Table 7**). Transcription factor motifs significantly enriched in DA-responsive genes included cyclic AMP response elements, which is consistent with previous reports demonstrating a key role for CREB in drug-induced transcriptional changes (*10, 37, 38*).

**Figure 2.**
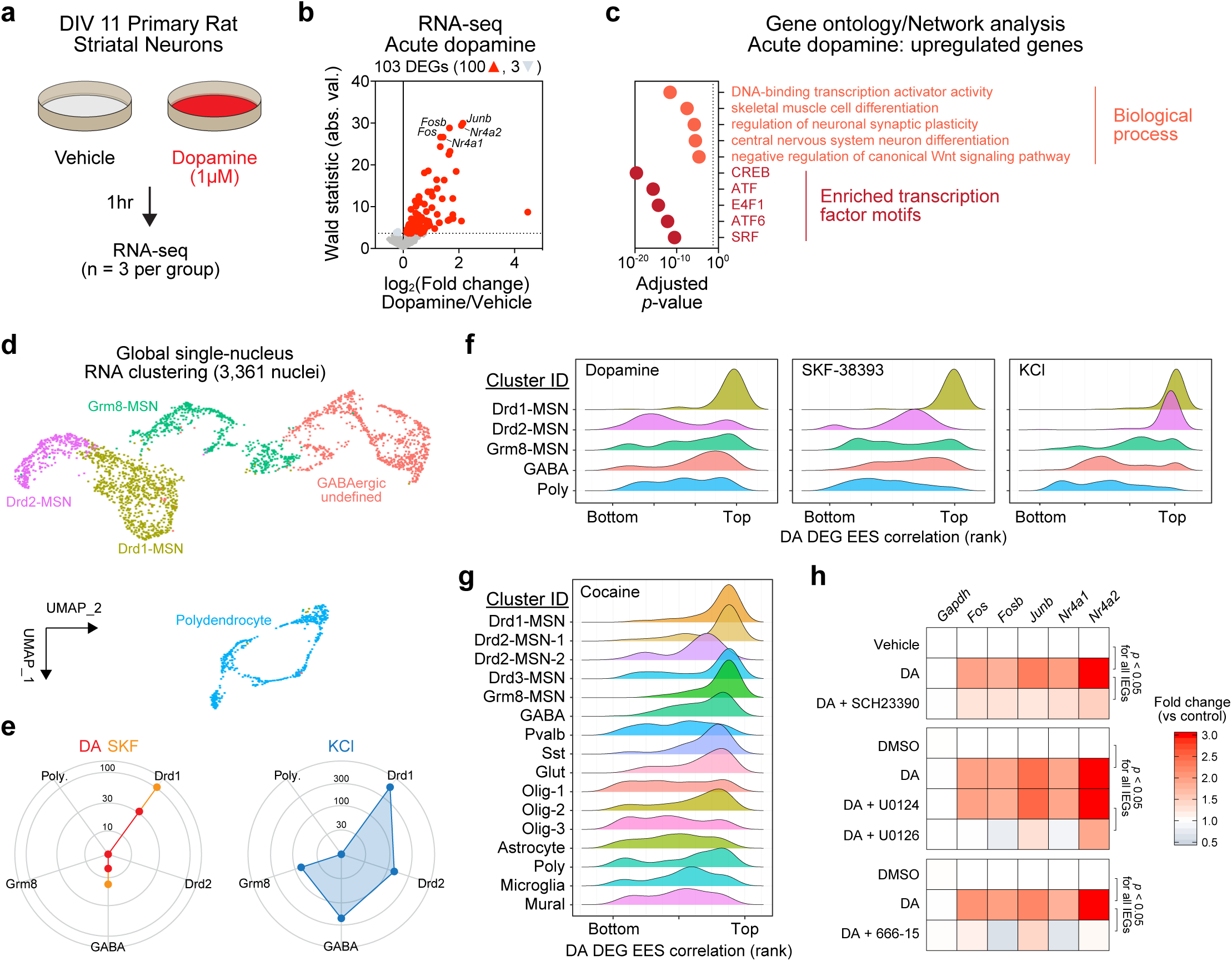
Identification of a cell-type specific dopamine-mediated transcriptional program. **a**, Illustration of the experimental setup for modeling dopamine receptor activation in medium spiny neuron cultures. **b**, Volcano plot showing signature of differentially expressed genes (DEGs) detected following 1 hr dopamine treatment. Dotted line on y-axis shows adjusted *p* value < 0.05. DESeq2 identified a total of 3 downregulated genes (light gray) and 100 upregulated genes (red), including immediate early genes (IEGs) *Fos, Fosb, Junb, Nr4a1*, and *Nr4a2*. **c**, Top 5 significant biological processes (gene ontology) and transcription factor binding site enrichment (motif analysis) for 100 upregulated genes in dopamine treated neurons. A maximum value was assigned to CREB (adjusted *p* value = 0). **d**, Single-nucleus RNA-seq from 3,361 nuclei identified 5 major cell classes in primary striatal neuron cultures. Neurons were treated with dopamine (50μM), the DRD1 agonist SKF-38393 (1μM), or depolarized with 25mM KCl. **e**, Differentially expressed genes by cell cluster. Dopamine and SKF-38393 induce a transcriptional response primarily in Drd1-MSNs, whereas KCl depolarization results in broad transcriptional alterations across neuronal classes. **f**, Unbiased signal identification (EES) identifies high correlation between DA-induced gene set (100 DEGs from bulk RNA-seq) and transcriptional response to dopamine or DRD1 agonist SKF specifically in Drd1 MSNs. Ridgeline plots show density of EES-gene correlation ranks for core DA signature genes (100 upregulated genes from 2b). This activation signature is present in Drd2 MSNs following depolarization, but not in response to DA. **g**, DA signature gene induction in multple MSN clusters following acute cocaine experience *in vivo* (clusters taken from Fig. 1b). **h**, RT-qPCR for representative DA transcriptional response genes following DRD1 receptor antagonist SCH-23390 (1μM), MEK inhibition (U0126, 1μM), or CREB inhibition (666-15, 1μM). All *p* values < 0.05 for DA IEGs.

To determine whether this core transcriptional response to DA occurs broadly across MSN subtypes or is limited to specific populations, we performed snRNA-seq from 3,361 cultured striatal neurons (mixed from male and female E18 rat brains) in 4 distinct treatment groups (Vehicle, 50µM DA, 1µM of the DRD1 receptor agonist SKF-38393, or 25mM potassium chloride (KCl) to induce neuronal depolarization). Cultured neurons were again largely divided into defined Drd1-MSNs, Drd2-MSNs, and Grm8-MSNs (**Fig. 2d**). Unlike scRNA-seq datasets from the adult NAc, primary striatal neuron cultures were devoid of oligodendrocyte, astrocyte, and microglia markers (due to the embryonic stage of tissue harvested for culture), but did harbor polydendrocytes positive for canonical marker genes *Pdgfra* and *Olig1* (**Supplementary Fig. 6**). Similar to cocaine experience *in vivo*, treatment with DA or SKF-38393 resulted in selective transcriptional activation of Drd1-MSNs (**Fig. 2e**). This activation included core DA signature genes identified from bulk RNA-seq (*Fos, Fosb, Junb, Nr4a1*, and *Nr4a2*), demonstrating robust induction of this gene program in Drd1-MSNs (**Supplementary Fig. 7; Supplementary Table 8**). Moreover, expression of nearly all of these genes was positively correlated with unbiased EES scores for DA and SKF-38393 treatments only in Drd1-MSNs (**Fig. 2f; Supplementary Table 9**), highlighting the key contribution of this program to overall transcriptional changes in these neurons. In contrast, both Drd1-MSN and Drd2-MSN clusters responded with massive transcriptional alterations following neuronal depolarization with KCl, which was reflected in the total number of DEGs, the fraction of cells activated, and the percentage of reads aligning to DA-responsive genes (**Fig. 2e** and **Supplementary Fig. 7**). Together, these results suggest that both Drd1- and Drd2-MSNs are capable of producing a robust transcriptional response involving the same core gene set, but that only Drd1-MSNs exhibit robust transcriptional responses to elevated DA neurotransmission in this model. Intriguingly, expression of this core gene set was positively correlated with EES score in the adult snRNA-seq dataset in multiple MSN clusters following cocaine experience (**Fig. 2g**), potentially reflective of multimodal (e.g., DA-dependent and DA-independent) contributions to this gene expression profile after cocaine exposure.

Given that DA-responsive genes were preferentially activated in Drd1-MSNs and that these genes are enriched for CREB binding motifs, we hypothesized that responsivity of these genes could be prevented via inhibition of the DRD1 receptor, interference with MAP kinase signaling cascades linked to CREB activation, or inhibition of CREB itself. To test these hypotheses, we performed RT-qPCR for key DA-response genes 1hr following DA stimulation (1µM) in the presence of either a DRD1 receptor antagonist (SCH-23390, 1µM), a Mitogen-activated protein kinase kinase (MEK) inhibitor (U0126, 1µM), or a CREB inhibitor (666-15, 1µM). In agreement with our prediction, we found that induction of each DA-responsive gene was inhibited by each of these treatments (**Fig. 2h**). These results confirm that DA-responsive gene programs are DRD1-dependent in this culture model system and demonstrate that these effects require MAP kinase signaling and classical CREB-mediated transcriptional mechanisms.

Although individual candidate genes in this DA-responsive transcriptional program have been shown to play key roles in drug-induced adaptations (*39, 40*), it is hypothesized that gene expression programs work in concert to exert downstream functional effects (*30, 41*). However, defining the significance of key gene expression programs has remained challenging due to the lack of tools capable of interrogating large-scale polygenic changes. Recent developments in CRISPR-based technology have enabled rapid and multiplexable transcriptional control in the mammalian central nervous system, providing an avenue to investigate how gene programs regulate normal and maladaptive brain states (*35, 42-44*). To simultaneously activate multiple genes in this DA-induced gene expression profile, we harnessed a neuron-optimized CRISPR activation (CRISPRa) system (**Fig. 3**) in which a catalytically dead Cas9 protein (dCas9) is fused to the strong hybrid transcriptional activator VPR (a concatemer of the herpes simplex viral protein VP16, the p65 subunit of NF-*κ*B, and the gammaherpesvirus transactivator Rta) (*35, 45*). This system allows multiplexed gene programming through the design of single guide RNAs (sgRNAs) that target the promoter region of selected genes. Using this system, we designed sgRNAs to target 16 of the genes most robustly altered by DA receptor activation (including *Fos, Fosb*, and *Nr4a1*; **Fig. 3a**). Following validation of each sgRNA using individual gene targeting in striatal cultures (**Supplementary Fig. 8**), we engineered custom multiplex sgRNA lentiviruses that contained eight sgRNAs each for a total of 16 unique sgRNAs (**Fig. 3a-d**). Expression of these vectors in striatal neurons in tandem with neuron-specific expression of the dCas9-VPR fusion protein enabled simultaneous induction of all genes (which we termed “Dopaplex”) as compared to a control multiplex virus expressing an sgRNA array targeting the bacterial gene *LacZ* (**Figure 3d**).

**Figure 3.**
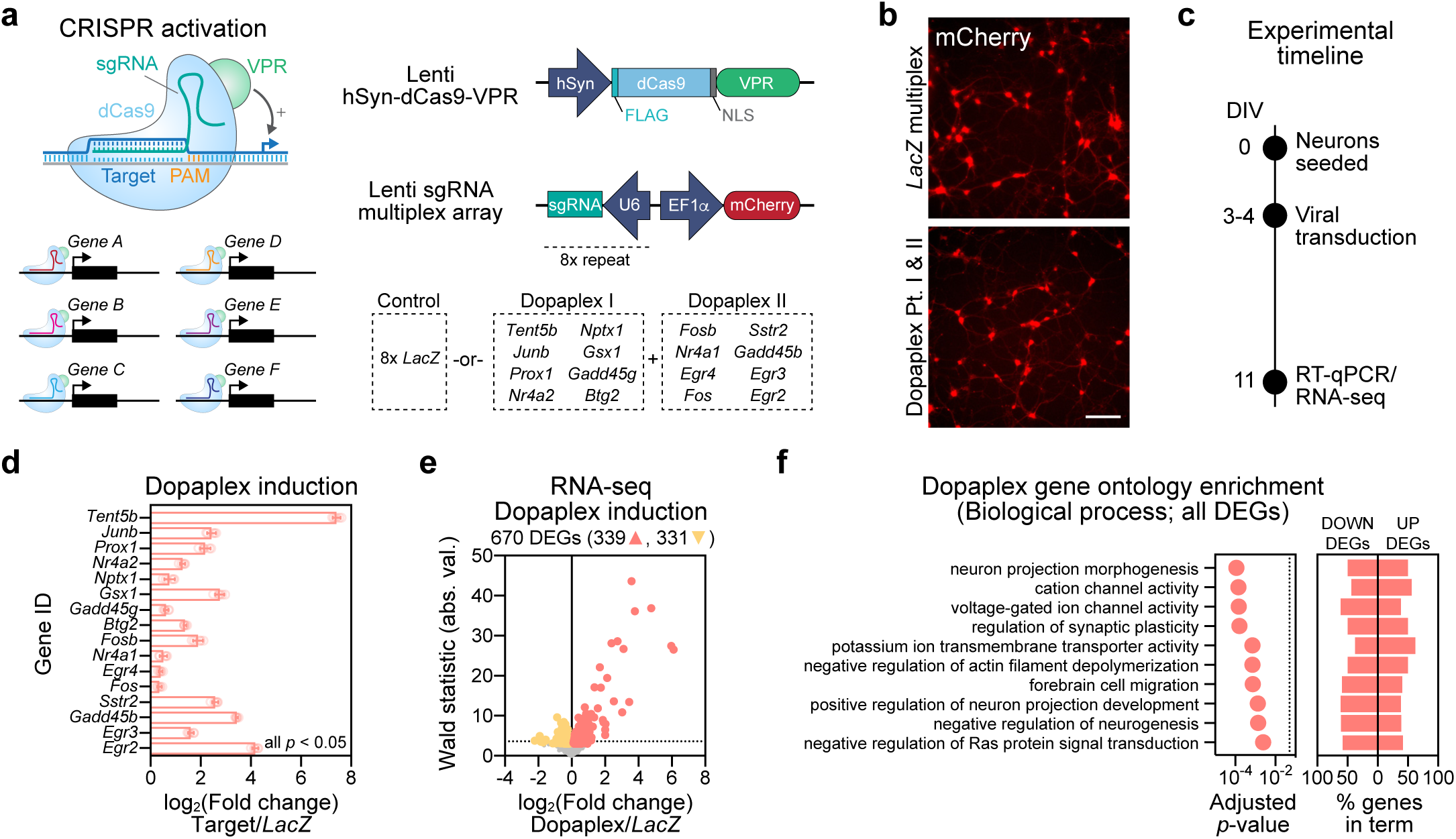
Multiplexed CRISPRa engineering to mimic dopamine-induced gene expression changes. **a**, Illustration of the CRISPRa multiplex vector approach expressing the dCas9-VPR activator fusion and multiplex single guide RNAs targeting either the bacterial *LacZ* gene (non-targeting control) or 16 of the top dopamine-induced genes (Dopaplex). **b**, Live cell imaging reveals successful transduction of the CRISPRa lentiviruses. Scale bar = 50µm. **c**, Experimental timeline for *in vitro* Dopaplex induction. Primary striatal neurons were generated and transduced with multiplexed CRISPRa constructs at DIV3-4. On DIV11, RNA was extracted and subjected to both RT-qPCR and RNA-sequencing to examine gene expression. **d**, dCas9-VPR increases gene expression of Dopaplex genes compared to the multiplexed *LacZ* control (*n* = 8, unpaired *t*-test; *Tent5b t*_*13.4*_ = 49.6, *p* < 0.0001; *Junb t*_*13.62*_ = 29.97, *p* < 0.0001; *Prox1 t*_*12.05*_ = 11.64, *p* < 0.0001; *Nr4a2 t*_*7.94*_ = 16.82, *p* < 0.0001; *Nptx1 t*_*13.75*_ = 7.722, *p* < 0.0001; *Gsx1 t*_*13.71*_ = 9.226, *p* < 0.0001; *Gadd45g t*_*13.98*_ = 9.435, *p* < 0.0001; *Btg2 t*_*9.586*_ = 29.71, *p* < 0.0001; *Fosb t*_*7.423*_ = 28.64, *p* < 0.0001; *Nr4a1 t*_*10.35*_= 29.52, *p* < 0.0001; *Egr4 t*_*8.382*_= 2.388, *p* = 0.0427; *Fos t*_*13.96*_ = 6.982, *p* < 0.0001; *Sstr2 t*_*9.132*_ = 20.70, *p* < 0.0001; *Gadd45b t*_*8.104*_ = 29.48, *p* < 0.0001; *Egr3 t*_*11.87*_ = 19.00, *p* < 0.0001; *Egr2 t*_*7.828*_ = 27.37, *p* < 0.0001;). All data are expressed as mean ± s.e.m. **e**, RNA-seq volcano plot showing DEGs detected by DESeq2 in Dopaplex verses *LacZ* multiplex targeting conditions. Upregulated genes (salmon, 339 genes) and downregulated genes (yellow, 331 genes) are indicated, and a standard Wald statistic cutoff corresponding to adjusted *p* < 0.05 is represented by the horizontal dotted line. **f**, Top 10 significant molecular function (GO) for all DEGs (excluding 16 Dopaplex genes) after Dopaplex induction.

Since many of the top DA-responsive genes were transcription factors, we next sought to define the transcriptional consequences of activating this program. RNA-seq comparisons following CRISPRa targeting *LacZ* or Dopaplex identified 670 DEGs, with 339 upregulated genes and 331 downregulated genes (**Figure 3e, Supplemental Table 10**). After removal of the 16 targeted Dopaplex genes from the gene list, gene ontology analysis revealed that critical neuronal processes were altered in the Dopaplex-targeted group, including genes involved in the regulation of ion channels, projection morphogenesis, and synaptic plasticity (**Figure 3f; Supplemental Table 11**). These results demonstrate that activation of a limited set of DA-sensitive genes can result in large-scale transcriptional consequences, including gene targets that might promote neurophysiological alterations.

Exposure to drugs of abuse or alterations in CREB signaling influence the intrinsic physiological properties of MSNs (*17, 19, 34*). Therefore, we sought to understand whether Dopaplex induction altered the physiological activity of MSNs using a high-throughput multielectrode array system in which striatal neurons are seeded directly on a 768 electrode array divided across 48 culture wells (**Fig. 4a-b**). Dopaplex-expressing neurons did not differ from non-targeting *LacZ* multiplexed control neurons in both the number of spontaneously active neurons and the mean action potential frequency (**Figure 4c-e**). However, action potential burst frequency was increased in Dopaplex-targeted neurons (**Fig. 4f-g**), demonstrating that this gene expression program alters MSN firing patterns. To determine if this phenotype could be mimicked by targeting individual genes in the Dopaplex program, we targeted CRISPRa machinery to either the transcription factor *Fosb* (which has been frequently implicated in cocaine action (*10, 39, 40*)), the nucleotidyltransferase *Tent5b* (the gene with the highest fold change after DA stimulation), or *LacZ* control. Neurons overexpressing either *Fosb* or *Tent5b* did not differ from *LacZ* sgRNA neurons in any physiological measure (active units, firing rate, or burst frequency; **Fig. 4h**), indicating that the increase in burst firing following induction of the Dopaplex program was not due to the induction of either of these genes alone.

**Figure 4.**
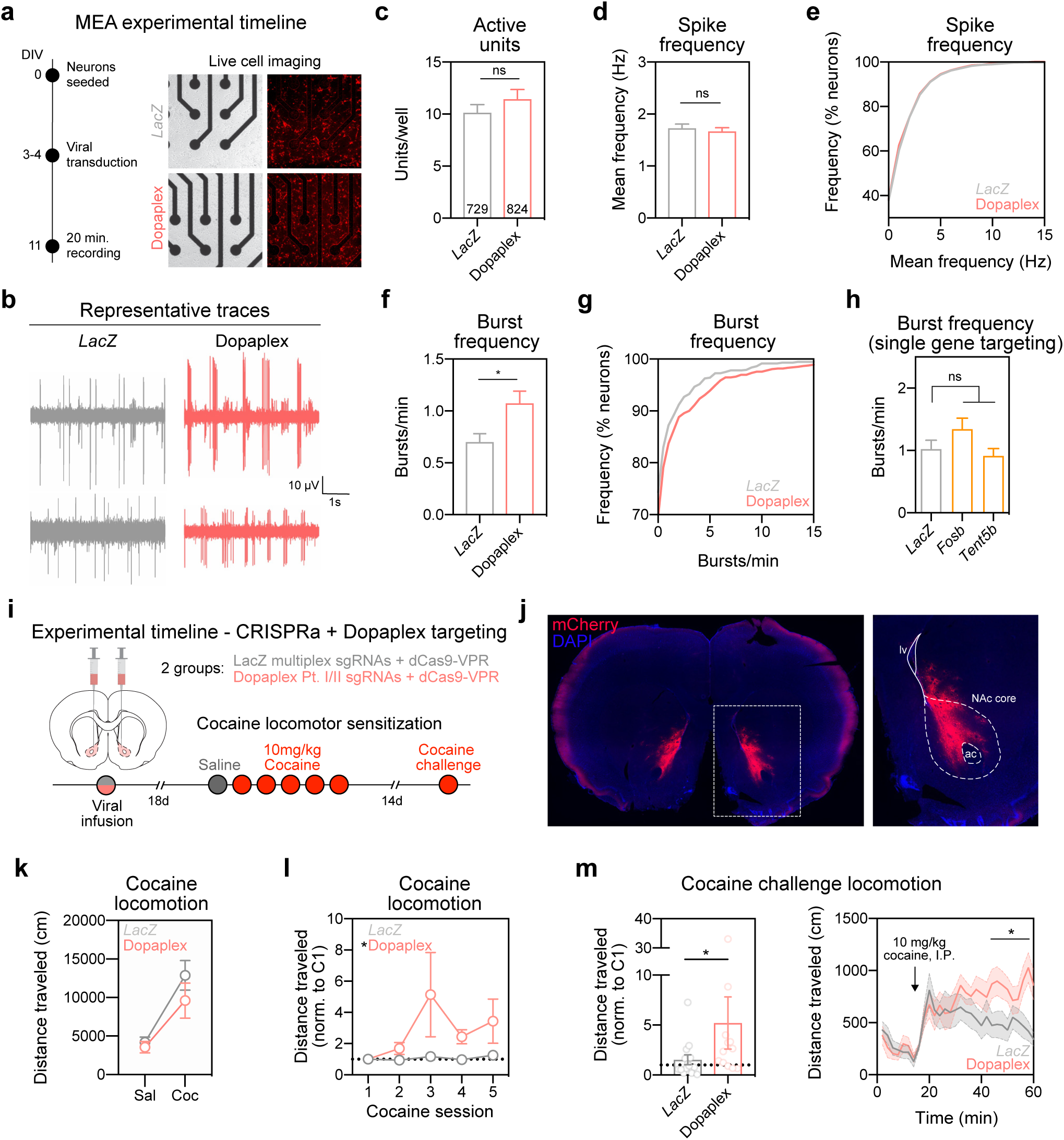
Dopaplex gene expression program increases bursting activity of MSNs and enhances cocaine locomotor sensitization. **a**, (left) Experimental timeline for viral transduction and MEA recordings. (right) Live cell imaging reveals successful transduction of the CRISPRa lentiviruses. **b**, Representative traces from 2 units each from the *LacZ* and Dopaplex targeting groups. The number of active units per well (**c**) and action potential frequency (**d-e**) did not change between *LacZ* and Dopaplex-targeted conditions (active units: *n* = 72 wells, Student’s *t*-test, *p* = 0.275; frequency: *n* = 729-824 neurons per group, Mann-Whitney *U* test, *U* = 296405, *p* = 0.6548). **f**, Burst frequency is increased in Dopaplex-targeted neurons (*n* = 729-824 neurons per group, Mann-Whitney *U* test, *U* = 282388, *p* = 0.0265). **g**, Cumulative distribution of burst frequency as a percent of all neurons. **h**, Burst frequency was not altered when CRISPRa was targeted to single genes within the Dopaplex gene expression program (n = 356-372 neurons per group, one-way ANOVA with Dunnett’s test for multiple comparisons, *F*_*2, 1087*_ = 2.295, *p* = 0.1012). **i**, Experimental timeline for viral transduction and cocaine locomotor sensitization. **j**, Representative image of sgRNA viral transduction (mCherry reporter) in the NAc. **k**, Absence of baseline locomotion difference between *LacZ* and Dopaplex targeted animals following saline injection or the first cocaine injection (*n* = 12-14 rats per group, Mann-Whitney *U* test, *U* = 65, *p* = 0.3474). **l**, Over 5 cocaine administration sessions, Dopaplex-targeted animals exhibited increased locomotion (*n* = 12-14 rats per group, two-way ANOVA with main effect of sgRNA, *F*_*1,24*_ = 4.412, *p* = 0.0464). **m**, Induction of sensitization with a cocaine challenge injection 14 days after the last cocaine pairing enhanced locomotion in Dopaplex-targeted rats (*n* = 12-14 rats per group, Mann-Whitney *U* test, *U* = 41, *p* = 0.0270). All data are expressed as mean ± s.e.m. Individual comparisons, \**p* < 0.05.

Our results suggest that DA signaling results in an altered transcriptional profile that supports increased burst firing of MSNs, which is consistent with reports that prior drug exposure alters the intrinsic properties of MSNs and enhances excitability (*17*). Therefore, we sought to understand how this gene expression program might regulate behavioral changes observed after exposure to drugs of abuse. One such behavior is locomotor sensitization, in which DA transmission in the NAc is potentiated as sensitization develops (*46, 47*). To examine the function of this Dopaplex gene expression program on cocaine locomotion sensitization, we stereotaxically infused CRISPRa lentiviral vectors targeting Dopaplex genes (or *LacZ* control) bilaterally into the NAc core (**Fig. 4i-j, Supplemental Figure 9**), a region critical for the development and maintenance of sensitization (*48, 49*). We previously validated robust and neuron-selective expression of this CRISPRa system *in vivo* within 2 weeks of viral delivery into the NAc, and demonstrated that this tool results in elevated protein levels of target genes (*35*). Eighteen days after viral infusion, rats underwent locomotor sensitization testing in a novel environment using a dose of cocaine (10mg/kg) that does not reliably produce robust sensitization (*50, 51*). Animals receiving the CRISPRa constructs targeting Dopaplex genes did not differ from *LacZ* multiplex controls in baseline locomotion after an injection of saline, nor in acute increases in locomotion following the first dose of cocaine (**Fig. 4k**). However, we observed a significant increase in the development of locomotor sensitization after 5 cocaine injections in Dopaplex-targeted rats (**Fig. 4l**). Likewise, this enhancement in locomotor activity was maintained 14 days after the last cocaine pairing by administration of a cocaine challenge dose (**Fig. 4m**). These results indicate that DA-mediated gene expression changes are sufficient to drive cocaine-related changes in neural physiology and behavior.

Following chronic drug administration, drug-induced dopaminergic signaling in the NAc is potentiated (*46, 47*), providing multiple opportunities for induction of long-lasting changes in gene expression that are observed after repeated drug use (*8, 9, 52*). While a candidate gene approach has been critical in understanding some of the fundamental transcriptional changes following exposure to drugs of abuse, activity-dependent transcriptional events produce coordinated changes that may be distinct from isolated changes in expression of a single gene. Our results shed new light on this process by identifying specific neuronal populations altered by cocaine experience, and further defining unique DA-responsive gene programs that control neuronal function and behavioral responses to cocaine. Together, these results allow comprehensive exploration of NAc transcriptional programs in an addiction-relevant model species, and identify novel target cells and genes for perturbation in the context of dopaminergic signaling. Given that addiction is a polygenic disorder that involves a complex interplay between genetic background and experience-dependent cellular responses, use of multiplexed gene regulation and cell-specific manipulations may enable a more complete understanding of how initial drug reinforcement transitions to long-term substance abuse disorders.

## Materials and Methods

### Animals

All experiments were performed in accordance with the University of Alabama at Birmingham Institutional Animal Care and Use Committee. Sprague-Dawley timed pregnant dams and 90-120-day-old male or female rats were purchased from Charles River Laboratories. Dams were individually housed until embryonic day 18 (E18) for cell culture harvest. Male or female adult rats were co-housed in pairs in plastic filtered cages with nesting enrichment in an AAALAC-approved animal care facility maintained between 23-24*°*C on a 12hr light/dark cycle with ad libitum food (Lab Diet Irradiated rat chow) and water. Bedding and enrichment were changed weekly by animal resources program staff. Animals were randomly assigned to experimental groups.

### Drugs

Cocaine hydrochloride was dissolved in sterile 0.9% sodium chloride. Drugs were purchased from Sigma-Aldrich (St. Louis, MO) and injected intraperitoneally (i.p.) at a dose of 20mg/kg or 10mg/kg for locomotor testing and locomotor sensitization, respectively. Cocaine solution was made fresh immediately before behavioral testing and protected from light. For *in vitro* experiments, drugs were diluted in Neurobasal medium (Invitrogen) immediately prior to treating cell culture plates. Dopamine hydrochloride (Sigma-Aldrich, H8502-5G) was dissolved in Neurobasal medium, and cells were treated at a dose of 1µM for RT-qPCR experiments or 50µM for snRNA-sequencing experiments. R(+)-SCH-23390 hydrochloride (Sigma-Aldrich, D054-5MG) and R(+)-SKF-38393 hydrochloride (Sigma-Aldrich, S101-5MG) were dissolved in sterile Milli-Q water, and cells were treated at a dose of 1µM. For MEK inhibitor experiments, U0124 (1µM; Millipore, 662006-1MG) and U0126 (1µM; Millipore, 662005-1MG) were dissolved in DMSO (Invitrogen, D12345). The CREB inhibitor 666-15 (1µM; Tocris, 5661), also called 3-(3-Aminopropoxy)-N-[2-[[3-[[(4-chloro-2-hydroxyphenyl)amino]carbonyl]-2-naphthalenyl]oxy]ethyl]-2-naphthalenecarboxamine hydrochloride, was dissolved in DMSO. Potassium chloride (Fisher Scientific, P330-500) was dissolved in Neurobasal media, and cells were treated with a 25mM dose.

### Neuronal Cell Cultures

Primary rat neuronal cultures were generated from E18 rat striatal tissue as described previously (*35,36*).Briefly, cell culture plates (Denville Scientific Inc.)were coated overnight with poly-L-lysine (Sigma-Aldrich; 50 µg/ ml), supplemented with 7.5 µg/mL laminin (Sigma-Aldrich), and rinsed with diH_2_O. Microelectrode arrays (MEAs; Axion Biosystems) were coated with polyethyleneimine (Sigma-Aldrich). Dissected striatal tissue was incubated with papain (Worthington LK003178) for 25 min at 37°C. After rinsing in complete Neurobasal media (supplemented with B27 and L-glutamine, Invitrogen), a single cell suspension was prepared by sequential trituration through large to small fire-polished Pasteur pipettes and filtered through a 100µm cell strainer (Fisher Scientific). Cells were pelleted, re-suspended in fresh media, counted, and seeded to a density of 125,000 cells per well on 24-well culture plates (65,000 cells/cm^2^) or 30,000 cells per well on 48-well MEA plates. Cells were grown in complete Neurobasal media for 11 days in vitro (DIV 11) in a humidified CO_2_ (5%) incubator at 37°C with half media changes at DIV 1, 4-5, and 8-9. MEAs received a one-half media change to BrainPhys (Stemcell Technologies Inc.) with SM1, L-glutamine supplements starting on DIV 4-5 and continued every 3-4 days. All MEA media was supplemented with penicillin-streptomycin.

### RNA extraction and RT-qPCR

Total RNA was extracted (RNAeasy kit, Qiagen) and reverse-transcribed (iScript cDNA Synthesis Kit, Bio-Rad). cDNA was subject to RT-qPCR for genes of interest, as described previously (*36*). A list of PCR primer sequences is provided in **Supplemental Table 12**.

### Tissue collection from adult NAc

One hour after i.p. injection of either saline vehicle or 20mg/kg cocaine, rats were euthanized by live decapitation, and the brain was rapidly removed and blocked into coronal sections in ice-cold Hibernate A media (Thermo Fisher, A1247501) supplemented with B27 and GlutaMax (Life Technologies, 35050-061). Tissue punches were then collected from coronal sections containing the NAc core and shell (∼ +1.68 AP from bregma; n=4 rats/treatment/sex) and rapidly frozen on dry ice. Tissue was stored at −80*°*C until the day of sequencing.

### Single Nuclei Dissociation

#### Frozen tissue from adult NAc

Frozen tissue punches were thawed slowly on ice prior to being chopped by scalpel 100 times in 2 orthogonal directions. Tissue from 4 animals in the same treatment condition was then combined and transferred to 5mL of chilled (4*°*C) Lysis Buffer (10mM Tris-HCl, 10mM NaCl, 3mM MgCl_2_ (Thermo Fisher, BP214-500), 0.1% Igepal in Nuclease-Free Water (Sigma, 18896-50ML)) for 15 min, mixing the tissue by inversion every 2 min. Lysis was quenched after 15 min with 5mL of complete Hibernate-A (Thermo Fisher, A1247501) supplemented with B27 and Glutamax (Life Technologies, 35050-061). Tissue was then triturated by fire-polished Pasteur pipette (2 pipettes of decreasing diameter; 8-10 passes per pipette) for 35 min and passed through a 40*μ*m pre-wet filter. The samples were then pelleted at 500rcf for 10 min at 4C followed by a wash in 10mL of Nuclei Wash & Resuspension Buffer (1X PBS, 1% BSA, 0.2U/ul NxGen RNase inhibitor (Lucigen, 30281-2)). Supernatant was then removed and the pellet was gently resuspended in a total volume of 800µl prior to 7-AAD (Thermo Fisher, 00-6993-50) staining and FACS to further purify the nuclei for sequencing. Immediately after FACS, nuclei were washed a final time at 200rcf in 10mL supplemented Hibernate-A containing 1% BSA for 10 min at 4*°*C to remove any remaining fine debris and nascent RNA. Nuclei were then brought to a concentration of 1,700 nuclei/ µL. Finally, an average total of 2,400 nuclei pooled from 4 rats per sex and treatment group were loaded into a single well of the Chromium Single Cell B Chip (10x Genomics catalog #1000074), utilizing 4 of the 8 available wells.

#### Striatal cell culture dissociation

On DIV11, striatal primary cells cultured on 12-well plates (250K/well) were treated with Neurobasal vehicle, 50µM DA, 1µM SKF, or 25mM KCL, and placed back into the incubator for 1hr. Cells were then washed with 1ml PBS per well, followed by a 5 min incubation period in 600µL Lysis Buffer. Cells were then gently pipette mixed 5 times with a P1000 before being transferred to a 1.5mL tube and centrifuged at 500rcf for 5 min at 4*°*C. Supernatant was then removed and the pellet resuspended in 100µL Nuclei Wash Buffer. Tubes for each treatment group were then combined, gently pipette mixed 5 times, and passed through a 40µM filter prior to FACS to further purify nuclei for sequencing. After FACS, 2,400 nuclei per treatment condition were loaded into individual wells of the Chromium Single Cell B Chip (10x Genomics, #1000074), utilizing 4 of the 8 available wells. Chromium single nuclei capture and barcoding were completed on the Chromium Single Cell Controller, according to manufacturer’s instructions.

### Single Nuclei RNA Sequencing and Analysis

Libraries were constructed according to manufacturer’s instructions using the Chromium Single Cell 3’ library construction kit (10X Genomics, #1000092), which utilizes version 3 chemistry for gene expression. 15,676 nuclei from adult rat NAc and 3,601 nuclei from cultured primary rat striatal neurons were sequenced on the Illumina NextSeq500 at the UAB Heflin Genomics Core to a depth of ∼38,000 reads per nuclei and ∼147,201 reads per nuclei, respectively. A Cell Ranger (v3.0.2) reference package was generated from the Ensembl Rn6 rat genome with a modified GTF file (version 95) to create a custom pre-mRNA package which ensured alignment to unspliced pre-mRNAs and mature mRNAs. Cell Ranger filtered outputs were analyzed with Seurat v3.0.2 using R v3.6.0(*53, 54*). Nuclei containing <200 features and >5% of reads mapping to the mitochondrial genome were removed from further analysis. Molecular count data from each GEM (Gelbeads in Emulsion) well was then log-normalized with a scaling factor of 10,000. Following normalization, and before dimensionality reduction, all GEM wells were integrated with FindIntegrationAnchors() and IntegrateData(), using 17 principal components (PCs) for the adult dataset and 10 PCs for the culture dataset. Dimensionality reduction is dependent on two main parameters, number of principal components and the resolution value. To determine the appropriate values, UMAPs were generated for every combination of 15 different PCs (2-30) and 20 different resolution values (0.1-2.0), using the standard Louvain algorithm. To produce UMAPs that capture previously defined cell types, we used 17 principal components and resolution values of 0.2 for the adult dataset, and 10 principal components with a 0.1 resolution value for the culture dataset. To validate that the clusters are specific and consist of known cell types, the log-normalized expression value of the top 2-6 genes from each striatal cell type, as identified by publicly available single-cell sequencing databases (e.g., DropViz.org (*23*) and mousebrain.org (*24*)), found within each dataset was overlaid on the UMAP using FeaturePlot(). To identify stimulation-dependent DEGs within each cell type, a Wilcoxon Ranked Sum test was performed on the log-normalized corrected unique molecular identifier (UMI) counts using custom R scripts based on Seurat source code. P-values were then adjusted using a Bonferroni correction based on the total number of genes identified within the dataset. Biological process gene ontology and transcription factor motif enrichment analysis for DEGs from specific cell clusters was performed in WEB-based Gene Set Analaysis Toolkit (WebGestalt) (*55*), using all protein coding genes as a reference set. Correction for multiple testing was performed using Benjamani-Hochberg adjustment, with statistical significance assessed at FDR < 0.05.

Enhanced Experimental Signal (EES) was calculated using the Manifold Enhancement of Latent Dimensions (MELD, version 0.0) (*29*) package on a cell-state graph generated using 100 PCA dimensions calculated with edge weights between cells generated through the Graphtools library with a knn=9 and a decay=10 using Python v3.7.3 after removing lowly expressed genes (genes not expressed in at least 10 cells (adult data set) or 5 cells (culture dataset)). The saline and cocaine experimental labels in the cell state graph were set with a smoothing parameter of Beta=1. The Markov Affinity-based Graph Imputation of Cells (MAGIC, v1.5.5)(*56*) package was utilized to impute gene expression using default settings on the cell-state graph. To determine genes which vary with the EES, pairwise Pearson correlation r values were calculated between each gene and the EES by cluster using SCPREP (v0.12.1). To generate the ridgeline plots, pairwise EES-gene Pearson correlation r values were calculated by cluster and ranked. Ridgeline plots reflect the ranks of 100 upregulated DEGs identified in the DA bulk RNA-seq experiment. The circos plot was generated using CIRCOS version 0.69 in Perl v5.18.4. The size of the outer rings represents the number of DEGs within each cluster. Genes were positioned by adjusted *p* values, and arcs were then drawn between genes that changed in the same direction in pairwise clusters and color coded to indicate neuron-neuon, glia-glia, and neuron-glia matches. ROC analysis was performed with a subset of snRNA-seq data from the parent Drd1-MSN cluster in Seurat v3.0.2. Dimensionality reduction was performed with 17 PCs and resolution of 0.2. For ROC evaluation, a binary classifier was built with each gene using FindMarkers(test.use = “roc”, assay = “RNA”, slot = “data”, logfc.threshold = 0, min.pct = 0.1).

### CRISPR-dCas9 construct design

CRISPRa experiments used lentivirus compatible plasmid constructs we previously optimized for robust neuronal expression (*35*). Gene-specific sgRNA targets were designed using online tools provided by the Zhang Lab at MIT (crispr.mit.edu) and CHOPCHOP (http://chopchop.cbu.uib.no/). To ensure specificity, all CRISPR RNA (crRNA) sequences were analyzed with National Center for Biotechnology Information’s (NCBI) Basic Local Alignment Search Tool (BLAST). A list of the target sequences is provided in **Supplemental Table 12**. crRNAs were annealed and ligated into the sgRNA scaffold using the BbsI cut sites. Plasmids were sequence-verified with Sanger sequencing.

Selected CRISPRa target genes were the genes with the highest confidence changes in gene expression following DA treatment. We applied a log_2_ fold change cutoff of > 1.2 on the bulk DA RNA-seq dataset, resulting in 23 potential target genes (**Supplemental Table 6**). We first designed sgRNAs to target each gene individually, excluding 3 genes due to significant predicted off-target localization of CRISPRa sgRNAs (**Supplemental Figure 2a**). At DIV3-4, primary striatal neurons were transduced with lentiviruses expressing sgRNAs targeting either a negative control sequence (the bacterial LacZ gene) or one of the high confidence DA-induced genes (**Supplemental Figure 8a**). Compared to the non-targeting control, we measured significant induction of 17 of the high confidence genes. To determine if these genes could be induced simultaneously, we pooled all 17 sgRNA viruses (**Supplemental Figure 8b**) and observed that all but one gene (*Fosl2*) was induced by this approach. Therefore, we excluded *Fosl2* from the multiplex CRISPRa targeting experiments. To deliver sgRNAs for each of these targets in a more efficient way, we generated new multiplex sgRNA lentiviruses that contained eight sgRNA sequences each (driven by distinct U6 promoters) for a total of 16 unique sgRNAs between 2 viruses (**Figure 3a**). The bacterial LacZ gene target was again used as a sgRNA non-targeting control by constructing an 8x LacZ sgRNA array. For multiplexing sgRNAs, individual sgRNAs targeting either LacZ or the DA-induced genes were inserted by PCR amplification and Golden Gate assembly into a custom-made destination plasmid co-expressing mCherry with unique BsmBI sites. All lentiviruses were produced as described previously (*35*), and titers for *in vitro* experiments were > 1×10^11^ GC/mL and > 1×10^12^ for *in vivo* experiments. Titers were determined with the Lenti-X RT-qPCR titer kit (Clonetech).

### Lentivirus production

Viruses were produced in a sterile environment subject to BSL-2 safety by transfecting HEK-293T cells (ATCC CRL-3216) with the specified CRISPR plasmid, the psPAX2 packaging plasmid, and the pCMV-VSV-G envelope plasmid (Addgene 12260 & 8454) with FuGene HD (Promega) for 40-48 hrs in supplemented Ultraculture media (L-glutamine, sodium pyruvate, and sodium bicarbonate) in a T225 culture flask. Supernatant was passed through a 0.45µm filter and centrifuged at 106,883rcf for 1 hr 45 min at 4°C. The viral pellet was resuspended in 1/100^th^ (*in vitro*) or 1/1000^th^ (*in vivo*) supernatant volume of sterile PBS and stored at −80°C. Physical viral titer was determined using Lenti-X qRT-PCR Titration Kit (Takara), and only viruses greater than 1×10^11^ GC/ml were used. Viruses were stored in sterile PBS at −80°C in single-use aliquots. For smaller scale virus preparation, each sgRNA plasmid was transfected in a 12-well culture plate as described above. After 40-48 hrs, lentiviruses were concentrated with Lenti-X concentrator (Takara), resuspended in sterile PBS, and used immediately.

### Multi Electrode Array Recordings

Single neuron electrophysiological activity was recorded using an Axion Maestro recording system (Axion Biosystems). E18 rat primary striatal neurons were seeded in 48-well MEAs at 30,000 cells/well, as described above. Each MEA well contains 16 extracellular recording electrodes and a ground electrode. Neurons were transduced with CRISPRa constructs on DIV 3-4 and MEA recordings were performed at DIV 11, while connected to a temperature- and CO_2_-controlled headstage (monitored at 37°C and 5% CO_2_). Electrical activity was measured by an interface board at 12.5 kHz, digitized, and transmitted to an external computer for data acquisition and analysis in Axion Navigator software (Axion Biosystems). All data were filtered using dual 0.01 Hz (high pass) and 5,000 Hz (low-pass) Butterworth filters. Action potential thresholds were set automatically using an adaptive threshold for each electrode (> 6 standard deviations from the electrode’s mean signal). Neuronal waveforms collected in Axion Navigator were exported to Offline Sorter (Plexon) for sorting of distinct waveforms corresponding to multiple units on one electrode channel, and confirmation of waveform isolation using principal component analysis, inter-spike intervals, and auto- or cross-correlograms. Further analysis of burst activity and firing rate was performed in NeuroExplorer (v.5.0).

### Bulk RNA-Sequencing

Bulk RNA-Sequencing (RNA-Seq) was carried out at Hudson Alpha Genome Services Laboratory or the Heflin Center for Genomic Science Genomics Core Laboratories at the University of Alabama at Birmingham. RNA was extracted, purified (RNeasy, Qiagen), and DNase-treated for three biological replicates per experimental condition. 1µg of total RNA underwent quality control (Bioanalyzer) and was prepared for directional RNA sequencing using NEBNext reagents (New England Biolabs) or SureSelect Strand Specific RNA Library Prep Kit (Agilent Technologies) according to manufacturer’s recommendations. PolyA+ RNA libraries underwent sequencing (50-75 bp paired-end directional reads; ∼25 M reads/sample) on an Illumina sequencing platform (HiSeq2000 or NextSeq2000).

### Bulk RNA-Seq Data Analysis

Paired-end FASTQ files were uploaded to the University of Alabama at Birmingham’s High Performance Computer cluster for custom bioinformatics analysis using a pipeline built with snakemake (*57*) (v5.1.4). Read quality, length, and composition were assessed using FastQC prior to trimming low quality bases (Phred < 20) and Illumina adapters (Trim_Galore! v04.5). Splice-aware alignment to the Rn6 Ensembl genome assembly (v90) was performed with STAR (*58*) v2.6.0c. An average of 88.4% of reads were uniquely mapped. Binary alignment map (BAM) files were merged and indexed with Samtools (v1.6). Gene-level counts were generated using the featureCounts (*59*) function in the Rsubread package (v1.26.1) in R (v3.4.1), with custom options (isGTFAnnotationFile = TRUE, useMetaFeatures = TRUE, isPairedEnd = TRUE, requireBothEndsMapped = TRUE, strandSpecific = 2, and autosort = TRUE). DESeq2 (*60*) (v 1.16.1) in R was used to perform count normalization and differential gene expression analysis with the application of Benjamini-Hochberg false discovery rate (FDR) for adjusted p-values. Differentially expressed genes (DEGs) were designated if they passed a p < 0.05 adjusted p-value cutoff and contained basemean > 39. Gene ontology (GO) analysis was conducted on DA (compared to vehicle control), and CRISPRa-induced Dopaplex gene expression program (compared to LacZ sgRNA control) using the ClueGO application in Cytoscape. Overrepresentation enrichment analysis was performed using non-redundant terms in biological process, using the protein-coding rat genome as a reference set. Enrichment analysis applied Benjamini-Hochberg correction for multiple comparisons and required a minimum of 3 genes per enriched GO term category. Transcription factor binding prediction used the WEB-based Gene Set Analaysis Toolkit (WebGestalt) (*55*).

### Stereotaxic Surgery

Naïve adult Sprague-Dawley rats were anaesthetized with 4% isoflurane and secured in a stereotaxic apparatus (Kopf Instruments). During surgical procedures, an anaesthetic plane was maintained with 1–2.5% isoflurane. Under aseptic conditions, guide holes were drilled using Paxinos and Watson(*61*) stereotaxic coordinates (all coordinates in respect to bregma: AP: +1.6 mm, ML: ±1.4 mm, DV: −7.0) to target the NAc core. All infusions were made using a gastight 30-gauge stainless steel injection needle (Hamilton Syringes) that extended into the infusion site. Bilateral lentivirus microinfusions of (1.5µl total volume per hemisphere) were made using a syringe pump (Harvard Apparatus) at a rate of 0.25µl/min. Injection needles remained in place for 10 min following infusion to allow for diffusion. Rats were infused bilaterally with either 1.5 µl of total lentivirus mix comprised of 0.75µl sgRNA and 0.75µl dCas9-VPR viruses in sterile PBS. After infusions, guide holes were covered with sterile bone wax and surgical incision sites were closed with nylon sutures. Animals received buprenorphine and carprofen for pain management and topical bacitracin to prevent infection at the incision site.

### Acute cocaine locomotion testing

For basic locomotor testing, naïve male (n=8) and female (n=8) animals were given a single i.p. injection of either cocaine (20mg/kg) or saline immediately before being placed in the activity chamber for 30 min of testing. Testing took place over two consecutive days between 2:00 PM and 5:30 PM, and sessions were counterbalanced across time by sex and treatment condition. The same female experimenter conducted locomotor testing and was present in the room during testing. Behavior chambers consisted of 43cm × 43cm Plexiglass locomotor activity chambers (Medical-Associates, St. Albans, VT) with opaque white wall covering and an open top. Each chamber included a 48-channel X-Y infrared array (Medical-Associates, St. Albans, VT) that was used to measure distance traveled in conjunction with Activity Monitor software (Medical-Associates, St. Albans, VT). Test chambers were cleaned with 0.0156% chlorhexidine and 70% ethanol at the beginning and end of each testing day, and cleaned with 70% ethanol in between trials. Animals were transported to the behavioral testing core 30 min prior to testing. Only the two animals currently undergoing locomotor testing were kept in the testing room; other animals were kept in an adjacent room. Behavior sessions were conducted during the light cycle, and the overhead lights and white noise generator in the behavior room remained on.

### Cocaine locomotor sensitization

After 2 weeks to allow for recovery and viral expression, animals were habituated to handling for a minimum of 4 days before locomotion testing began. Locomotor sensitization experiment sessions were 1 hr, with a 15 min habituation time before an i.p. injection of either saline (day 1) or 10mg/kg cocaine (days 3, 5, 7, 9, 11, and 25).

### Immunohistochemistry

Immunohistochemistry (IHC) was performed as described previously(*35*). Adult male rats were transcardially perfused with formalin (1:10 dilution in PBS, Fisher) 24-48 hr after the cocaine challenge. Brains were removed and post-fixed for 48 hr in formalin, then sliced at 50µm using a vibratome. Tissue slices were permeabilized with 0.25% Triton X-100 in PBS, then blocked for 1 hr at room temperature with blocking buffer (1x PBS with 10% Thermo Blocker BSA and 1% goat serum). Expression of mCherry was examined by incubation with an anti-mCherry primary antibody (1:500 in PBS with 10% Thermo Blocker BSA and 1% goat serum, Abcam catalog #ab183628) overnight at 4°C. Slices were washed three times with PBS and incubated for 1 hr at room temperature with a fluorescent secondary antibody (Alexa Fluor 546 goat anti-rabbit, Thermo Fisher Scientific catalog #A-11010, 1:500). Slices were washed three times with PBS and mounted onto microscope slides with Prolong Gold anti-fade medium (Invitrogen) containing 4,6-diamidino-2-phenylindole (DAPI) stain as a marker for cell nuclei. 4X images of the viral infusion site were taken on a Nikon TiS inverted fluorescent microscope and viral placement was mapped onto to coronal diagrams adapted from Paxinos and Watson (*61*) (**Supplementary Fig. 9**). Animals with strong viral expression in at least one hemisphere were included in behavioral studies, and one animal was excluded due to targeting.

### Statistical Analysis

Sample sizes were calculated using a freely available calculator (Lenth, R. V. (2006-9). Java Applets for Power and Sample Size [Computer software]. Available at http://www.stat.uiowa.edu/~rlenth/Power). Correspondence between the DA and SKF datasets was determined with a linear regression. Transcriptional differences from RT-qPCR experiments were compared with an unpaired t-test with Welch’s correction or one-way ANOVA with Dunnett’s or Tukey’s post-hoc tests where appropriate. MEA data was compared with Mann-Whitney U-tests or a one-way ANOVA. Sensitization data was compared with either an unpaired t-test with Welch’s correction or two-way ANOVA. Statistical significance was designated at *α* = 0.05 for all analyses. Statistical and graphical analyses were performed with Prism software (GraphPad). Statistical assumptions (e.g., normality and homogeneity for parametric tests) were formally tested and examined via boxplots.

## Data Availability

Sequencing data that support the findings of this study are available in Gene Expression Omnibus (GSE137568, GSE137658, GSE137759, and GSE137763). All relevant data that support the findings of this study are available by request from the corresponding author (J.J.D.). With the exception of sgRNA plasmids, CRISPRa constructs are available in the Addgene plasmid repository.

## Acknowledgements

This work was supported by NIH grants DA039650, DA034681, and MH114990 (J.J.D.), DA042514 (K.E.S.), DA041778 (F.A.S.). L.I. is supported by the Civitan International Research Center at UAB. Additional assistance to J.J.D. was provided by the UAB Pittman Scholars Program and the UAB Flow Cytometry Core. We thank Samantha Black for assistance with stereotaxic surgery and all current and former Day Lab members for assistance and support. We specifically thank Mikael Guzman Karlsson and Svitlana Bach for their contributions to establishment of single-cell RNA sequencing platforms in the lab. We thank the UAB Flow Cytometry Core for assistance with 10X snRNA-seq and FACS.

## Author contributions

J.J.D., J.J.T., C.G.D., M.E.Z., and R.A.P. conceived of single nuclei RNA-seq experiments. M.E.Z. and J.J.T. performed behavioral assays and generated tissue for single-nucleus RNA-seq, with assistance from R.A.P. and C.G.D. C.G.D. and J.J.T developed the single-nuclei dissociation protocol with assistance from R.A.P., M.E.Z., and the UAB Flow Cytometry Core. R.A.P., C.G.D., & J.J.D. performed statistical and graphical analysis from single-cell RNA-seq datasets with assistance from L.I. and J.J.T.. M.E.Z. performed DRD1 antagonist, MEK inhibitor, and CREB inhibitor experiments. K.E.S. and J.J.D. conceived of the CRISPR/ dCas9 multiplexing experiments. K.E.S., L.I., R.A.P., and J.J.D. performed statistical and graphical analysis of bulk RNA-seq datasets. K.E.S. designed, cloned, and validated custom sgRNA arrays and performed electrophysiological and behavioral assays for CRISPR/dCas9 multiplexing, with assistance from A.J.B., S.T., F.A.S., and N.A.G. All projects were supervised by J.J.D.. K.E.S. and J.J.D. wrote the main text of the manuscript. K.E.S., C.G.D., J.J.T., R.A.P., M.E.Z., and J.J.D. wrote the methods section of the manuscript. All authors have approved the final version of the manuscript.

## Competing Interests

The authors declare no competing financial interests.

---

**Supplementary Figure 1.**
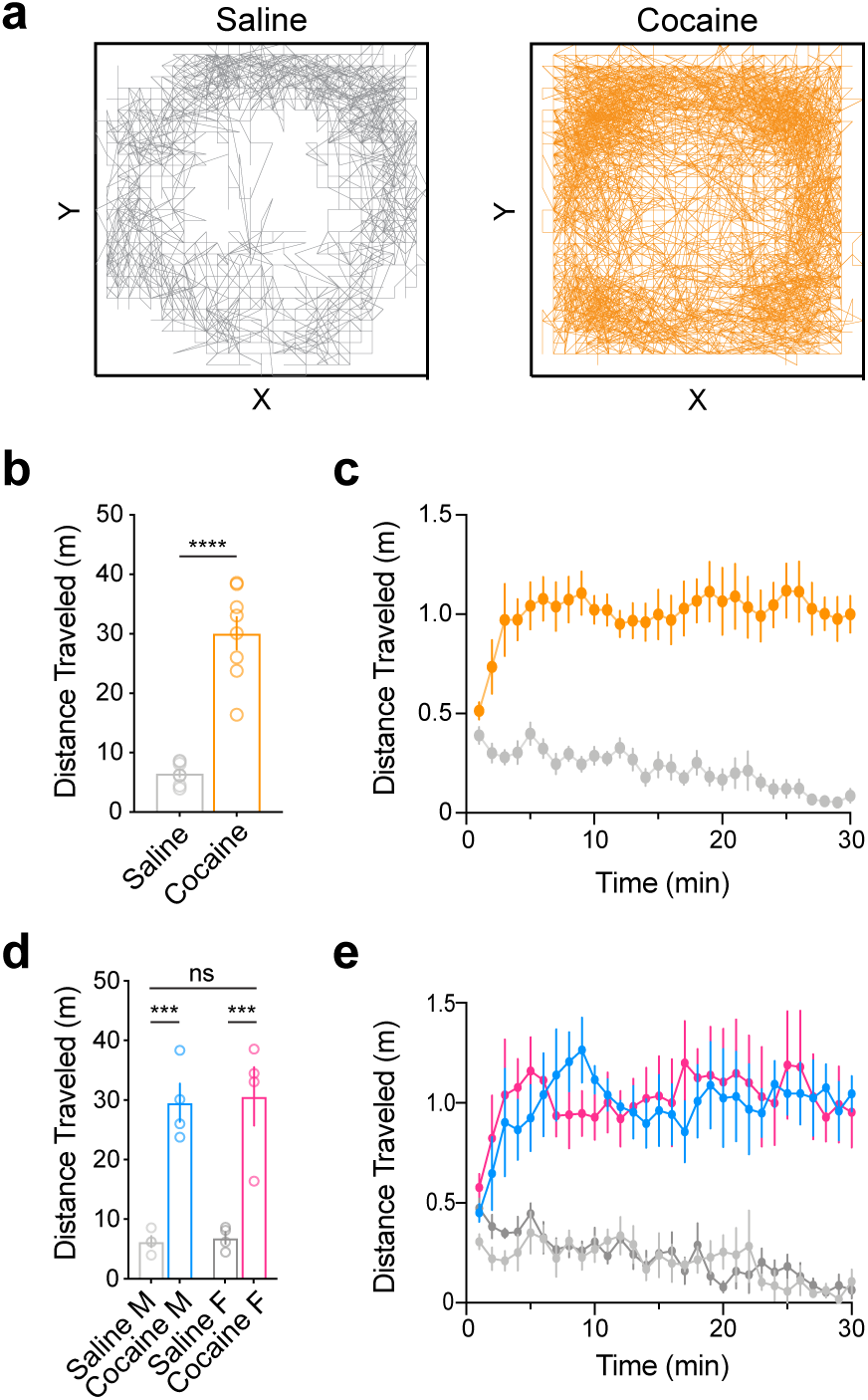
Cocaine experience significantly increases locomotor activity in both male and female rats. **a**, Representative activity trace of saline (n = 8) and cocaine (n = 8) animals over the 30 minute test session. **b**, Mean total distance traveled is significantly greater in rats who received a single i.p. injection of cocaine (20 mg/kg, n=8) compared to saline controls (n = 8; unpaired t test, *t*_*(14)*_ = 8.45, *p* < 0.0001, *R*^*2*^ = 0.84). **c**, Locomotor activity is increased in the cocaine group (20 mg/kg, n = 8) across all timepoints, compared to saline controls (n = 8). **d**, While there was a siginificant effect of cocaine on locomotor activity (20 mg/kg; one-way ANOVA *F*_*(3,12)*_ = 20.55, *p* < 0.0001, *R*^*2*^ = 0.84), Tukey’s *post hoc* analysis revealed no sex differences in the mean distance traveled in response to saline (n = 4/sex; *p* = 0.9988) or acute cocaine (n = 4/sex; *p* = 0.9940). **e**, Within each treatment group, there are no sex differences in locomotor activity at any timepoint (n = 4/sex/treatment group). All data are expressed as mean ± s.e.m. \*\*\**p* < 0.001, \*\*\*\**p* < 0.0001. M = male (light grey, blue), F = female (charcoal, pink).

**Supplementary Figure 2.**
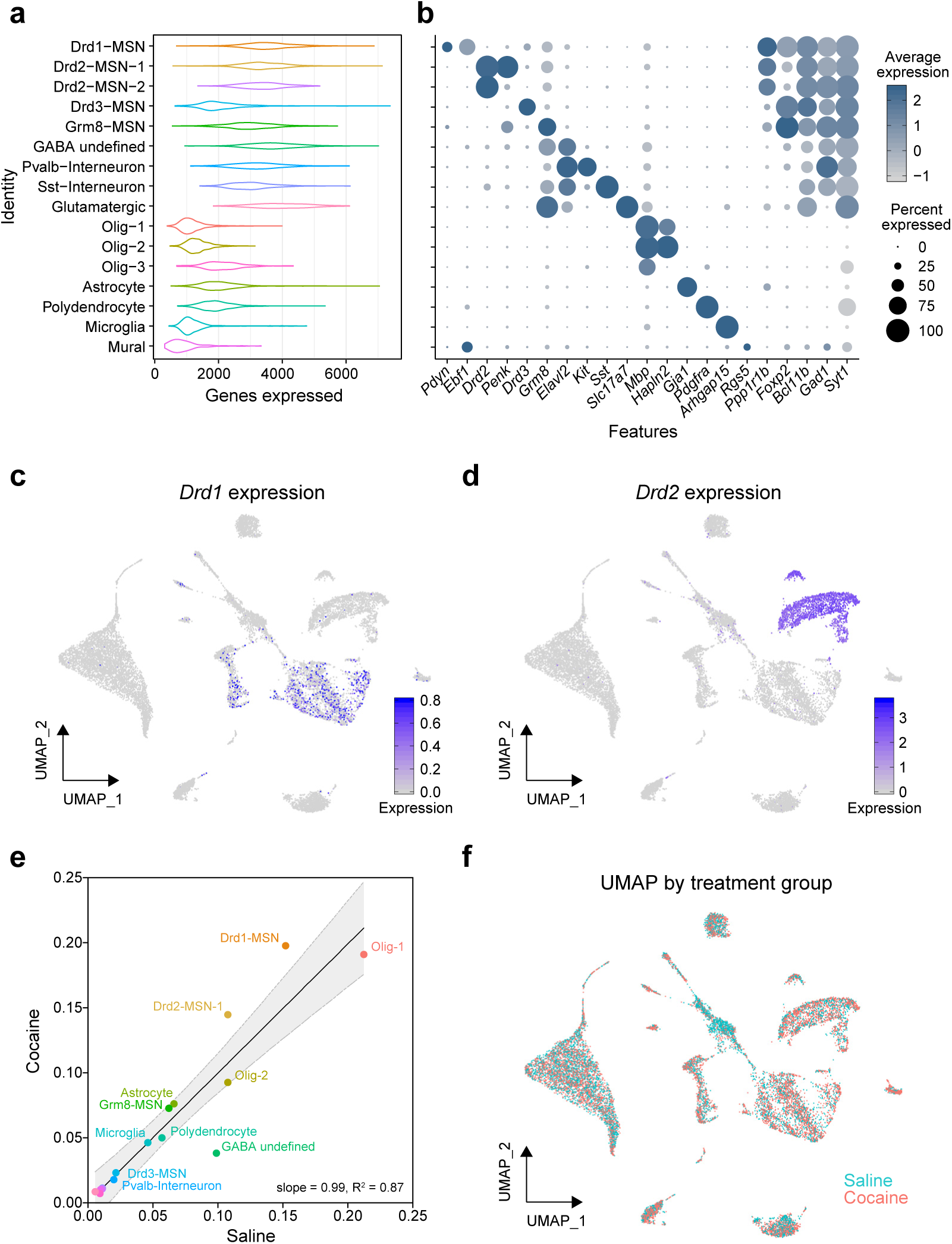
Identification of cell types within the rat nucleus accumbens. **a**, Violin plots indicating the distribution of number of total genes expressed by cell class. **b**, Dot plot indicating the average expression and percent of cells expressing marker genes of each identified celltype. **c-d**, Enrichment of *Drd1* and *Drd2* transcripts within identified clusters. **e**, Fraction of cocaine and saline treated neurons within each cluster. **f**, UMAP showing distribution of cells from saline and cocaine-treated rats.

**Supplementary Figure 3.**
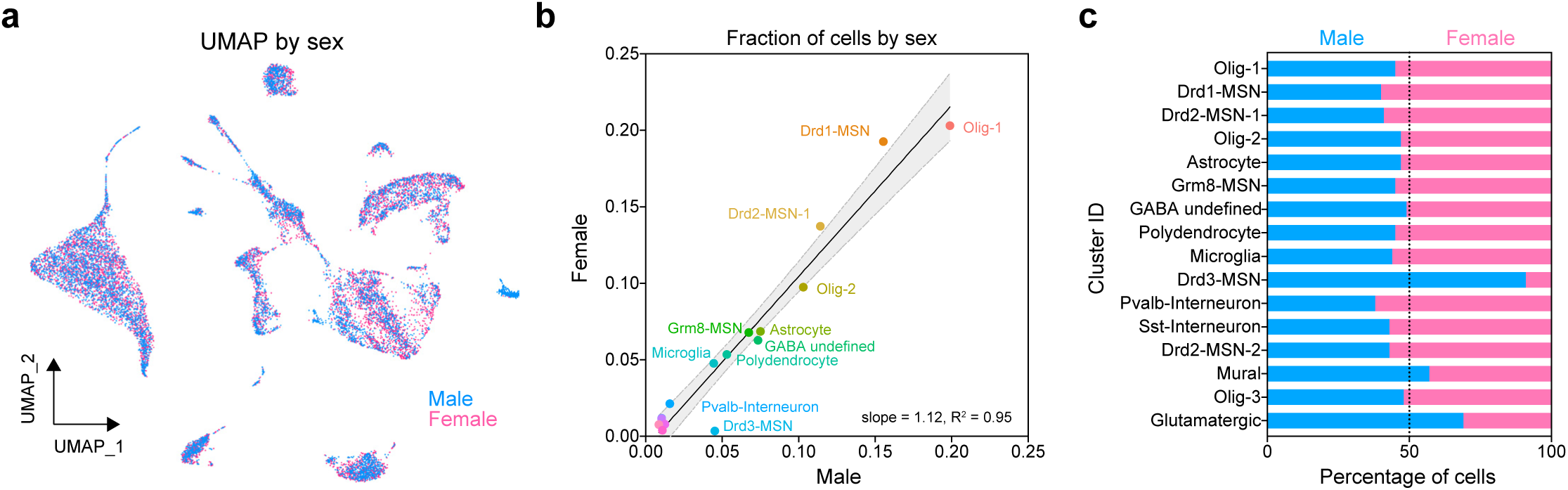
Sex differences in NAc cell distribution. **a**, Global UMAP showing male and female cells following Seurat integration. **b**, Linear regression of male and female cell fractions reveals highly similar cell distribution in nearly all cell clusters, with the exception of Drd3-MSNs. **c**, Percentage of each cluster accounted for by male and female cells. Over 90% of Drd3-MSNs were obtained from male animals.

**Supplementary Figure 4.**
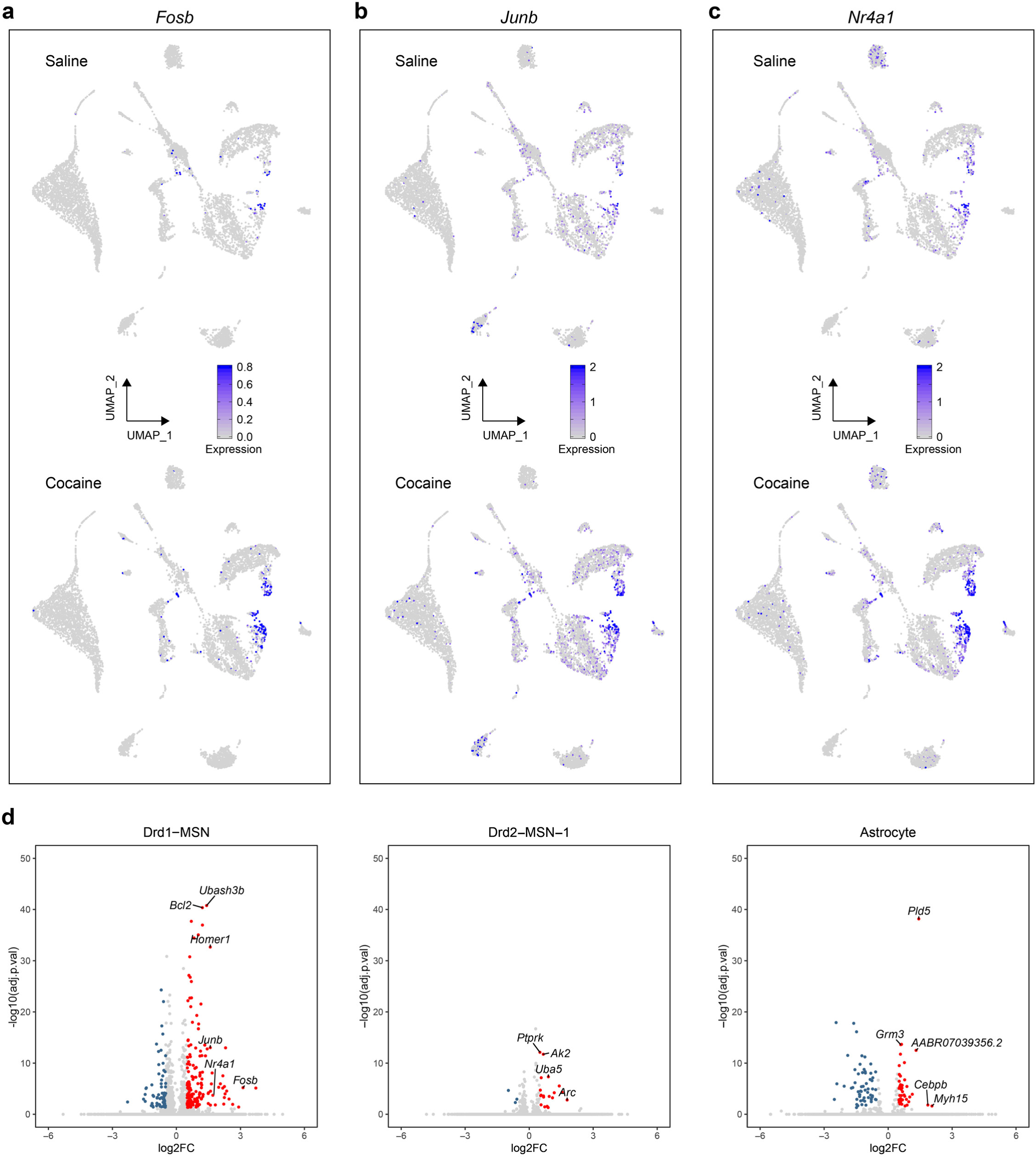
Cocaine induces immediate early gene response in Drd1-MSNs. **a-c**, Global UMAPs plotting expression values of *Fosb, Junb*, and *Nr4a1.* For each gene, cocaine increases mRNA abundance in the Drd1-MSN population, **d**, Volcano plots showing all DEGs (adjusted *p* value < 0.05, absolute value of log2(fold change) > 0.5) for three distinct cell clusters.

**Supplementary Figure 5.**
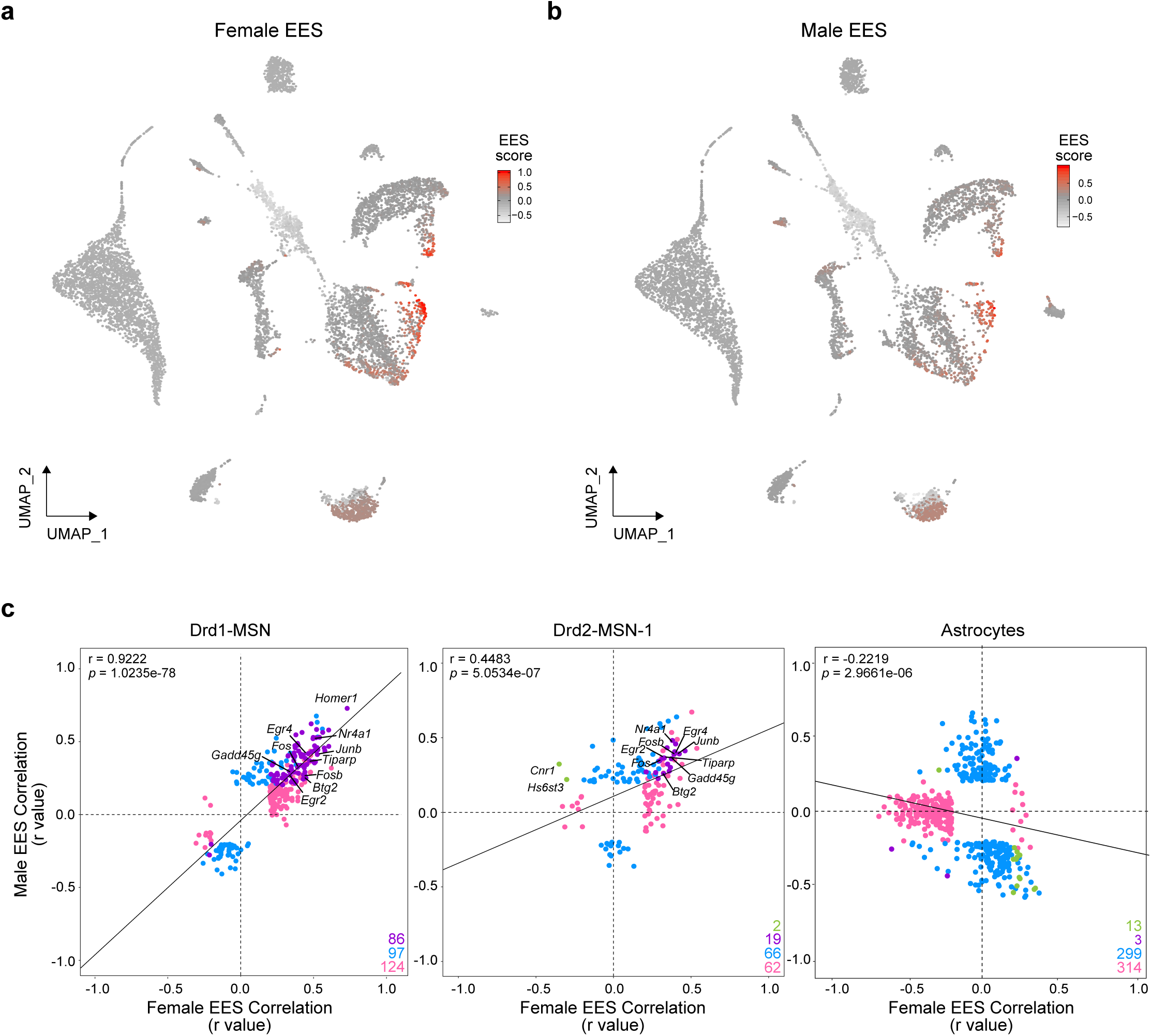
Contribution of sex to cocaine-mediated transcriptional responses in specific cell types in adult rat NAc. **a-b**, UMAPs showing enhanced experimental signal (EES) generated independently for each sex. **b**, EES-gene correlation correspondence plots reveal highly similar cocaine-induced transcriptional responses in Drd1- and Drd2-MSN clusters. In contrast, cocaine-mediated transcriptional responses in astrocytes were divergent in males and females.

**Supplementary Figure 6.**
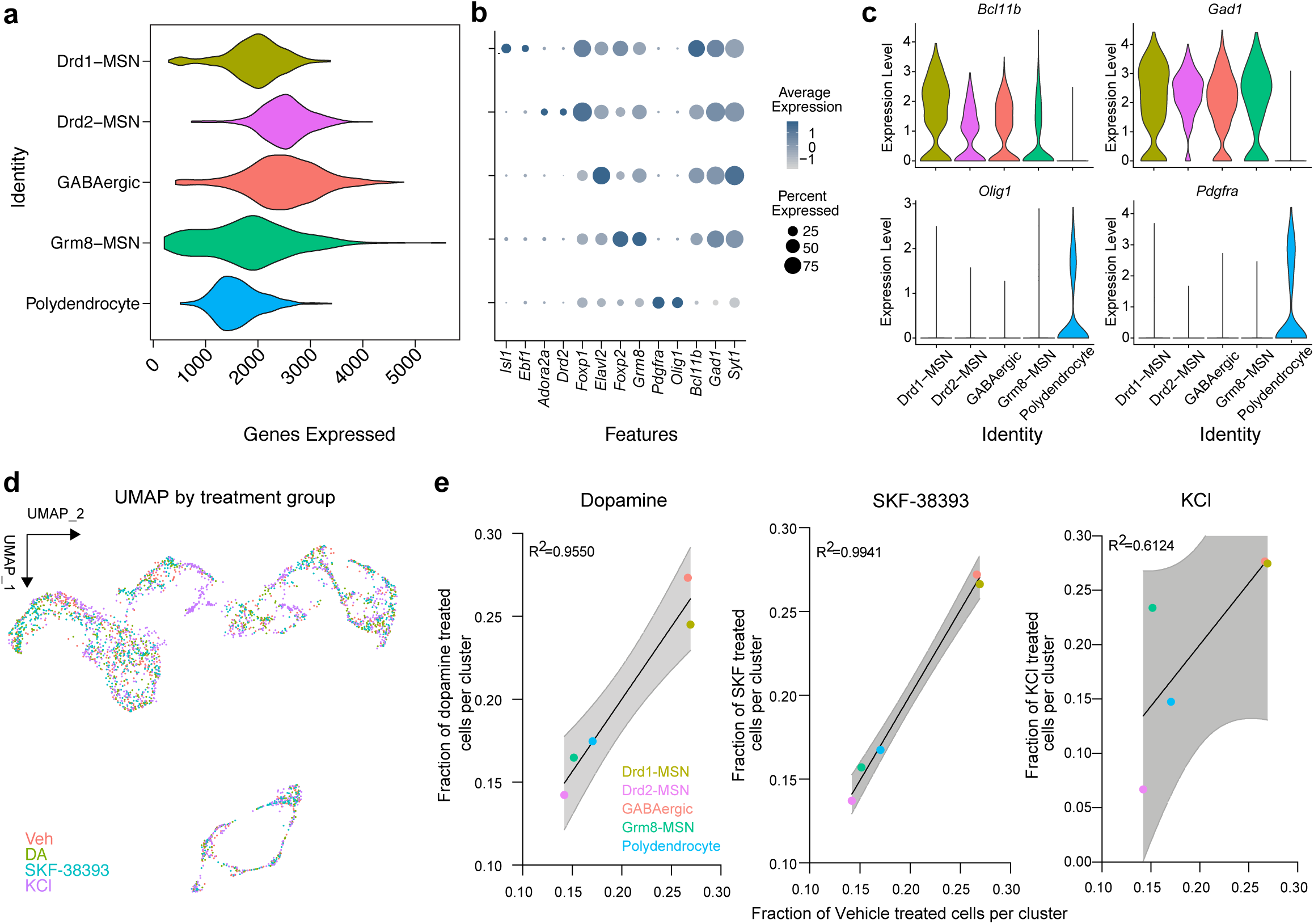
Identification of cell types within rat embryonic striatal neurons. **a**, Violin plots indicating the distribution of the number of genes expressed in each cluster **b**, Dot plot indicating the average expression and percent of cells expressing marker genes of each identified celltype. **c**,Violin plots indicating the distribution of neuronal (*Bcl11b,Gad1*) and polydendrocyte (*Olig1, Pdgfra*) marker genes. **d**, Distribution of dopamine, KCl, or SKF treated cells **e**, Fraction of dopamine, KCl, or SKF treated cells treated neurons within each cluster.

**Supplementary Figure 7.**
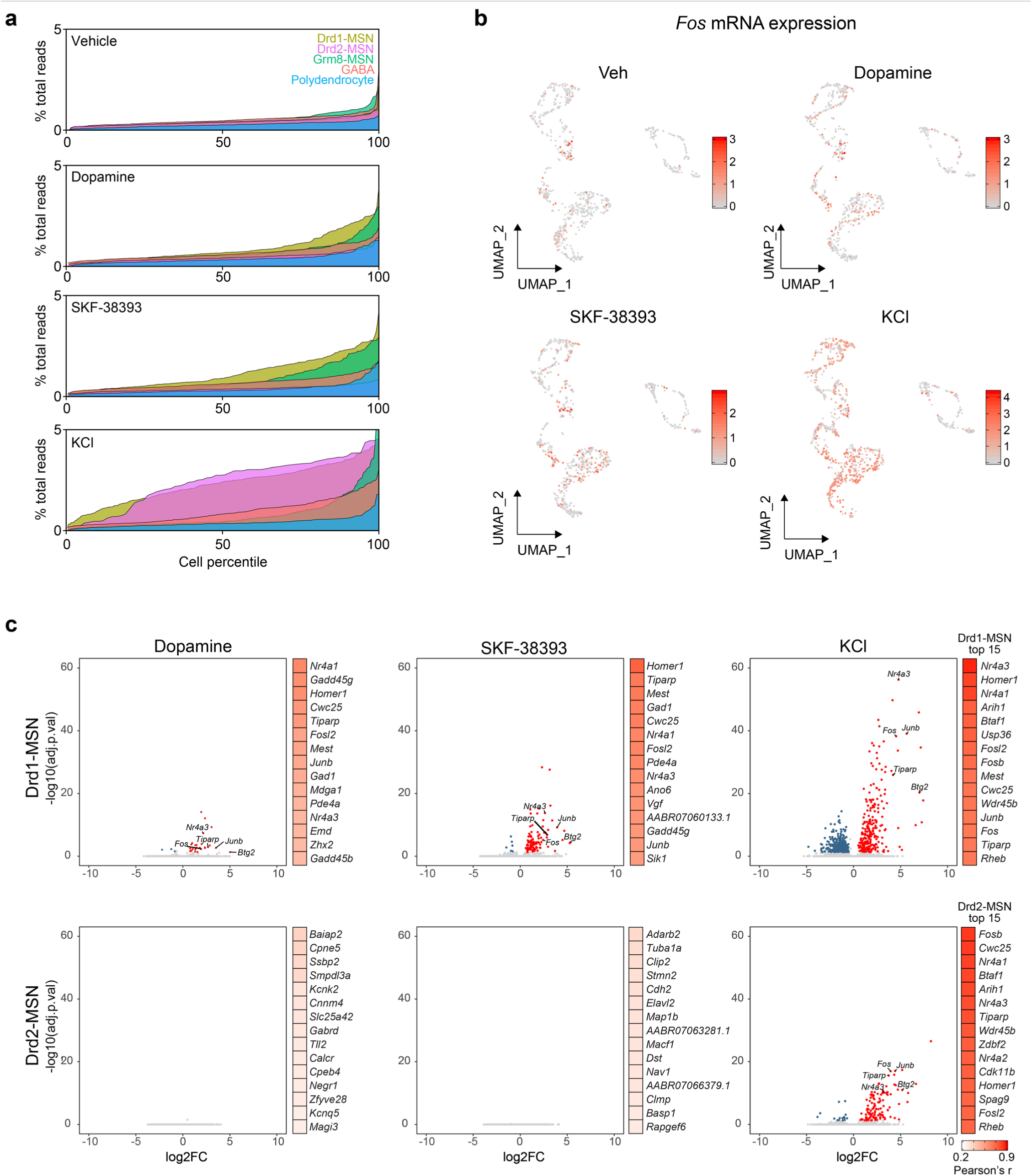
Figure S7 Treatment-induced changes in gene expression in cultured striatal neurons are cell-type specific. **a**, Percent of reads aligning to 100 DA-DEGs for each cell across treatment groups from cell culture snRNA-seq experiment Cells percentiled due to different numbers in each duster, **b**. UMAPs of Fos gene expression changes across dopamine. SKF, and KCI treatment groups, **c**. Volcano plots showing all DEGs (adjusted *p* value < 0.05, absolute value of log2(fold change) > 0 5) and top 15 EES correlating genes for Drd1- and Drd2-MSNs Dopamine and SKF-38393 induce transcnptional responses in Drd1-MSNs only, while KCI induces a large-scale transcnptional response in both cell types.

**Supplementary Figure 8.**
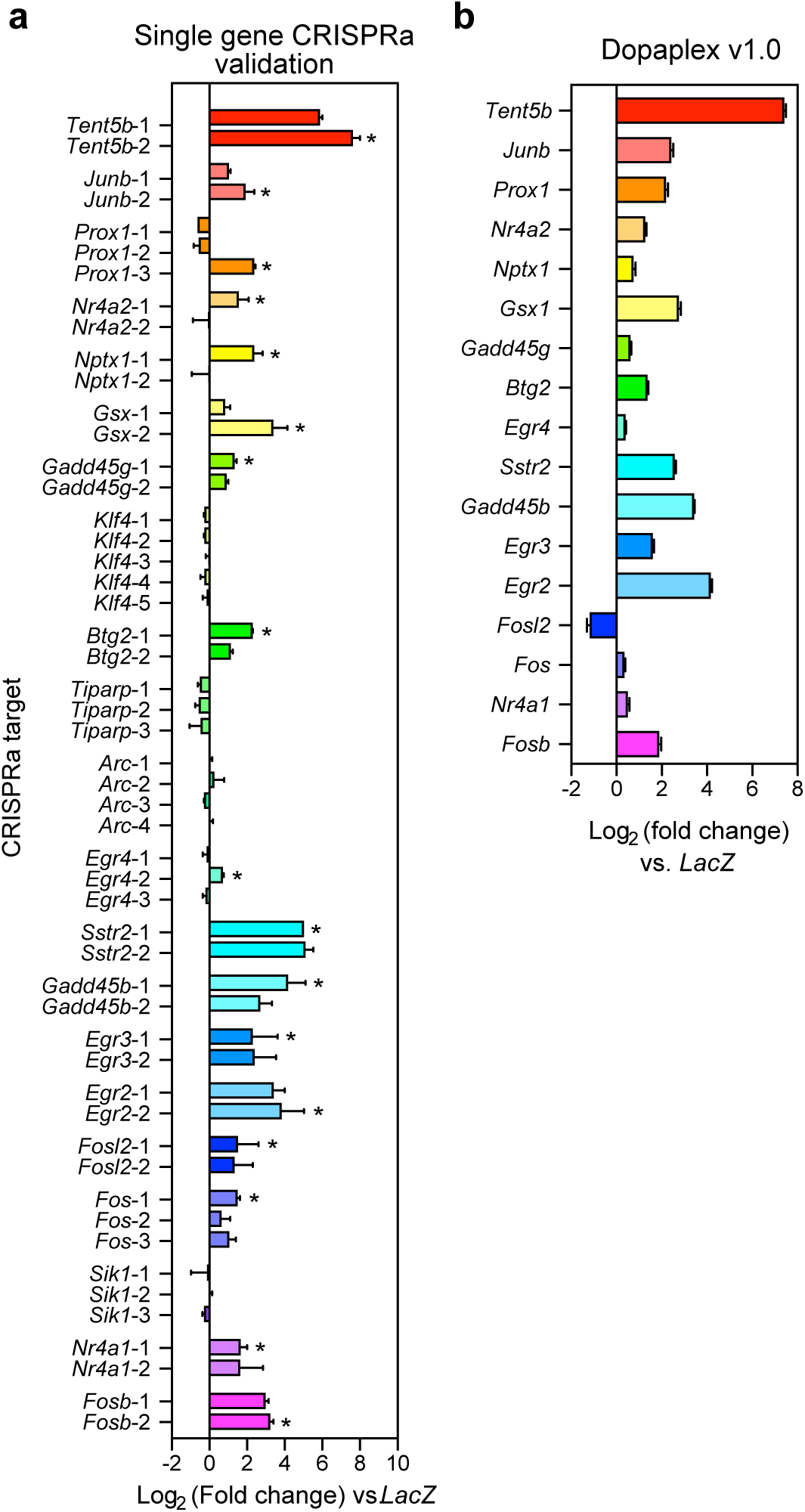
CRISPRa validation at single gene targets. **a**, Single guide RNAs targeted with dCas9-VPR to DA-induced genes results in validated induction in 16 out of 23 genes, as compared to a *LacZ* control sgRNA (*n* = 2 per group). Asterisks indicate sgRNAs selected for multiplexed targeting. (**b**) Lentiviral vectors expressing sgRNAs validated individually to upregulate target genes were pooled to target 17 DA-induced genes simultaneously. Simultaneous targeting resulted in induction of all but one gene (*n* = 5 per group). All data are expressed as mean ± s.e.m.

**Supplementary Figure 9.**
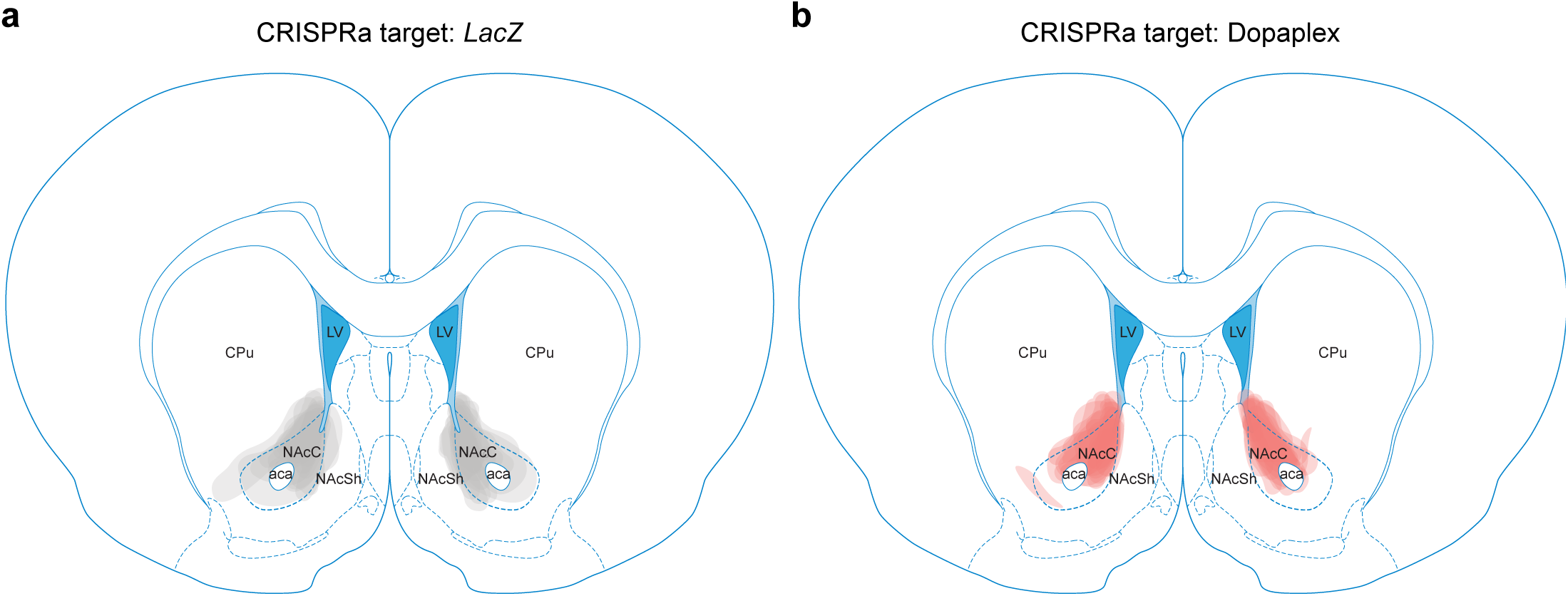
Viral placement validation for animals infused with CRISPRa constructs. CRISPRa lentivirus targeted to either the bacterial *LacZ* (**a**) or Dopaplex (**b**) were infused into the nucleus accumbens core (NAcC). Schematics of target regions are adapted from Paxinos and Watson. NAcC: nucleus accumbens core; NAcSh: nucleus accumbens shell; CPu: caudate putamen; LV: lateral ventricle.

